# Fatal neuroinvasion and SARS-CoV-2 tropism in K18-hACE2 mice is partially independent on hACE2 expression

**DOI:** 10.1101/2021.01.13.425144

**Authors:** Mariano Carossino, Paige Montanaro, Aoife O’Connell, Devin Kenney, Hans Gertje, Kyle A. Grosz, Maria Ericsson, Bertrand R Huber, Susanna A. Kurnick, Saravanan Subramaniam, Thomas A. Kirkland, Joel R. Walker, Kevin P. Francis, Alexander D. Klose, Neal Paragas, Markus Bosmann, Mohsan Saeed, Udeni B. R. Balasuriya, Florian Douam, Nicholas A. Crossland

**Author notes:** Co-corresponding author’s contact information: Florian Douam, PhD; Nicholas Crossland, DVM. These authors contributed equally to the work (Paige Montanaro, Aoife O’Connell, Devin Kenney, Florian Douam, Nicholas A. Crossland).

## Abstract

Animal models recapitulating distinctive features of severe COVID-19 are critical to enhance our understanding of SARS-CoV-2 pathogenesis. Transgenic mice expressing human angiotensin-converting enzyme 2 (hACE2) under the cytokeratin 18 promoter (K18-hACE2) represent a lethal model of SARS-CoV-2 infection. The precise mechanisms of lethality in this mouse model remain unclear. Here, we evaluated the spatiotemporal dynamics of SARS-CoV-2 infection for up to 14 days post-infection. Despite infection and moderate pneumonia, rapid clinical decline or death of mice was invariably associated with viral neuroinvasion and direct neuronal injury (including brain and spinal neurons). Neuroinvasion was observed as early as 4 dpi, with virus initially restricted to the olfactory bulb supporting axonal transport via the olfactory neuroepithelium as the earliest portal of entry. No evidence of viremia was detected suggesting neuroinvasion occurs independently of entry across the blood brain barrier. SARS-CoV-2 tropism was not restricted to ACE2-expressing cells (e.g., AT1 pneumocytes), and some ACE2-positive lineages were not associated with the presence of viral antigen (e.g., bronchiolar epithelium and brain capillaries). Detectable ACE2 expression was not observed in neurons, supporting overexpression of ACE2 in the nasal passages and neuroepithelium as more likely determinants of neuroinvasion in the K18-hACE2 model. Although our work incites caution in the utility of the K18-hACE2 model to study global aspects of SARS-CoV-2 pathogenesis, it underscores this model as a unique platform for exploring the mechanisms of SARS-CoV-2 neuropathogenesis that may have clinical relevance acknowledging the growing body of evidence that suggests COVID-19 may result in long-standing neurologic consequences.

**IMPORTANCE:** COVID-19 is predominantly a respiratory disease caused by SARS-CoV-2 that has infected more than 191 million people with over 4 million fatalities (2021-07-20). The development of animal models recapitulating distinctive features of severe COVID-19 is critical to enhancing our understanding of SARS-CoV-2 pathogenesis and in the evaluation of vaccine and therapeutic efficacy. Transgenic mice expressing human angiotensin-converting enzyme 2 (hACE2) under the cytokeratin 18 promoter (K18-hACE2) represent a lethal model of SARS-CoV-2 infection. Here, we show lethality of this model is invariably associated with viral neuroinvasion linked with viral replication and assembly. Importantly, pneumonia albeit invariably present was generally moderate with the absence of culturable infectious virus at peak neuroinvasion. The dynamics of viral neuroinvasion and pneumonia were only partially dependent on hACE2. Overall, this study provides an in-depth sequential characterization of the K18-hACE2 model following SARS-CoV-2 infection, highlighting its significance to further study the mechanisms of SARS-CoV-2 neuropathogenesis.

## INTRODUCTION

The world is experiencing the devastating effects of the Coronavirus Disease 2019 (COVID-19) pandemic, a highly contagious viral respiratory disease caused by the newly emerged betacoronavirus, Severe Acute Respiratory Syndrome Coronavirus-2 (SARS-CoV-2) (1–3). The initial index case was reported at a seafood market in Wuhan, Hubei Province, China in late 2019 (1). While still under investigation, it has been postulated that the progenitor of SARS-CoV-2 may have originated from horseshoe bats (*Rhinolophus affinis*) or Malayan pangolins (*Manis javanica*) that, following spill over into humans, acquired the genomic features leading to adaptation and human-to-human transmission (1). SARS-CoV-2 has a high transmissibility rate, and, to date, it has infected nearly 194 million people, resulting in over 4 million fatalities (2021-07-28) (4). COVID-19 causes respiratory disease of variable severity, ranging from mild to severe, with the development of acute respiratory distress syndrome requiring intensive care and mechanical ventilation (3, 5–7). Numerous comorbidities including hypertension, obesity, and diabetes, among others, are affiliated with an increased risk of developing severe COVID-19 (5, 6, 8–10). Furthermore, a proportion of infected patients go on to develop poorly understood neurological signs and/or symptoms mostly associated with the loss of smell and taste (anosmia and ageusia), headache, dizziness, encephalopathy (delirium), and ischemic injury (stroke), in addition to a range of less common symptoms (5, 7, 11–17). Multiple studies have identified either SARS-CoV-2 RNA and/or protein in the brain of COVID-19 patients with the olfactory neuroepithelium postulated as a portal of entry (18, 19). COVID-19 has severely challenged health care systems around the globe, with the urgent need for medical countermeasures including the development of efficacious vaccines and therapeutics.

Animal models permissive to SARS-CoV-2 that could serve as suitable models to help better understand the pathogenesis of COVID-19, while simultaneously assisting in the development and evaluation of novel vaccines and therapeutics to combat this disease, are critically needed (20–22). While various animal models (mice, hamsters, non-human primates, ferrets, minks, dogs, and cats) have been evaluated to date (22–30), none faithfully recapitulates all the pathological features of COVID-19. The main limitation in the development of suitable murine models of COVID-19 is related to the virus entry mechanism: SARS-CoV-2 binds to target cells via interaction between the viral spike protein (S) and the host angiotensin-converting enzyme 2 (ACE2), considered to be the major host entry receptor (31). The low binding affinity between the S protein and murine ACE2 (mACE2) renders conventional mouse strains naturally resistant to infection, posing a challenge in the development of murine models of COVID-19 (31–34). These difficulties have been circumvented by the development of transgenic murine models that express human ACE2 (hACE2) under different promoters including hepatocyte nuclear factor-3/forkhead homologue 4 (HFH4), and cytokeratin 18 (K18) (30, 35–38). The transgenic murine model expressing hACE2 under a K18 promoter (namely K18-hACE2) was developed by McCray et al in 2007 to study SARS-CoV-1 (36), which shares the same host receptor as SARS-CoV-2 (39).

SARS-CoV-2 infection of K18-hACE2 mice results in up to 100% lethality, analogous to that reported for SARS-CoV (30, 36, 38). Early reports communicated lethality to be associated primarily with severe lung inflammation and impaired respiratory function, suggesting that this model can recapitulate features of the respiratory disease observed in severe cases of COVID-19 (30, 38). However, the confounding impact of neuroinvasion and its role in the clinical decline of SARS-CoV-2 infected K18-hACE2 mice is becoming more readily acknowledged (19, 40, 41).

Under K18 regulation, the expression of hACE2 is reported to be limited mainly to airway epithelial cells and enterocytes lining the colonic mucosa, to a lower degree within kidney, liver, spleen, and small intestine, and to a relatively minor level of expression in the brain (36). However, the cellular distribution of ACE2, and particularly hACE2, in tissues of K18- hACE2 mice remains largely undetermined. We hypothesized that the nature, severity, and outcome of disease in K18-hACE2 mouse model is not solely dictated by the expression and tissue distribution of hACE2 and that increased lethality in this model is ultimately related to neuroinvasion, in part driven by regional ACE2 overexpression in the nasal passages promoting retrograde axonal transport through the olfactory nerve. To investigate this hypothesis, we undertook a comprehensive spatiotemporal analysis of histologic and ultrastructural changes, cellular distribution of viral protein and RNA, viral loads, and antibody titers, along with a detailed analysis of the distribution of *hACE2* mRNA, ACE2 protein, and its correlation with SARS-CoV-2 tropism as it pertains to this model.

Although SARS-CoV-2 protein and RNA were detected in ACE2-expressing cells such as olfactory neuroepithelium (ONE) and alveolar type 2 (AT2) cells, we found that SARS-CoV-2 primarily infected neurons and alveolar type 1 (AT1) pneumocytes, which lacked detectable ACE2 protein. Our results support neuroinvasion as the primary cause of mortality in the K18-hACE2 mouse model. This claim was supported by the observation that viral load and titers peaked in the brain when animals began to meet euthanasia criteria or succumb to disease. This was clinically reflected by onset of profound hypothermia and onset of neurological signs including tremors. Neurons in terminal animals display prominent spongiotic degeneration and necrosis with concurrent detection of abundant viral protein, RNA, and virus particle assembly. Although pneumonia was uniformly observed and peaks at 7 dpi, it was of moderate severity with declining viral loads compared to that of the brain. We found that several histologic hallmarks of severe COVID-19 were lacking in this model, (i.e., lack of diffuse alveolar damage and microthrombi), suggesting pneumonia plays a contributing role rather than the primary determinant of lethality in this model. Interestingly, the absence of detectable ACE2 expression in neurons suggested that viral neuroinvasion is a mechanism partially independent of ACE2.

Altogether, this study expands the current knowledge on the K18-hACE2 murine model to study severe COVID-19. Our findings will help refine utilization of this model for providing a relevant understanding of the molecular mechanisms driving neuropathogenesis and pulmonary pathology.

## RESULTS

### SARS-CoV-2 is invariably fatal in infected K18-hACE2 mice with evidence of neuroinvasion

K18-hACE2 mice inoculated intranasally with SARS-CoV-2 (1×10^6^ plaque-forming units [PFU]; n=35 [n=19 male and n=16 female) began losing weight as early as 4 days post-infection (dpi) irrespective of sex, with maximum weight loss occurring at 6-7 dpi (18.4% □ 6.8% in male mice, 21.4% ± 1.8% in female, and combined 19.7% ± 5.2%; Fig. 1A). Trends in weight loss paralleled increasing clinical scores and declines in core body temperature, with the latter two precipitously increasing or decreasing respectively near the time of death (Fig. 1B, C). SARS-CoV-2-infected K18-hACE2 mice exhibited neurological signs starting 6 dpi, characterized by profound stupor, tremors, proprioceptive defects, and abnormal gait, with most animals euthanized or found dead in their cage at 6 and 7 dpi (∼94%; 33/35 [Fig. 1D]). At the time of death (6-7 dpi), the median clinical score was 3 (interquartile range = 1) and the mean body temperature was 30.9 ± 3.0 °C. Two male mice survived to the end of the 14-day observation period and did not display hypothermia during the course of the observation period, a feature that was consistently observed in animals that succumbed to disease or met euthanasia criteria.

**Fig. 1.**
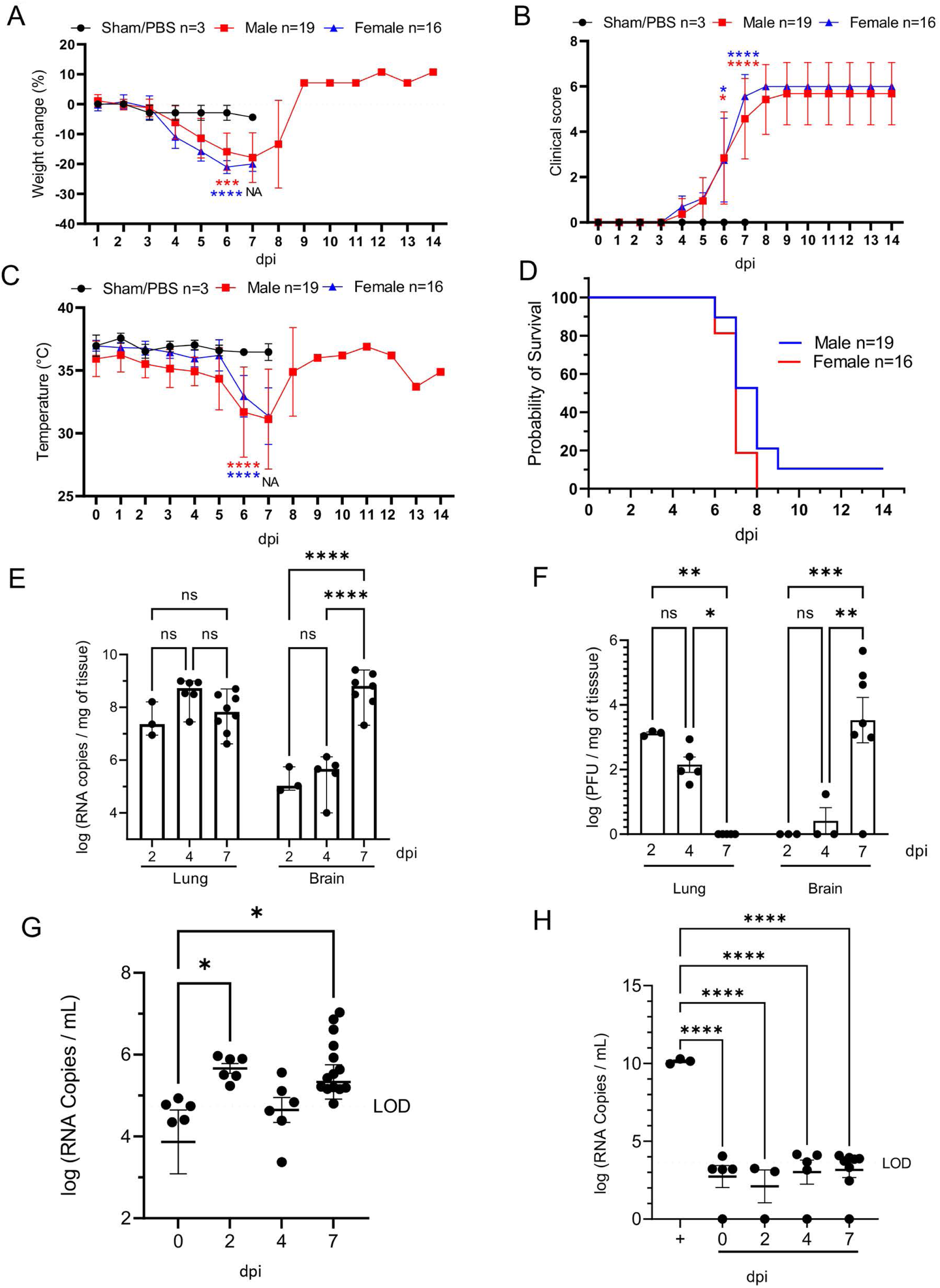
SARS-CoV-2 caused lethal disease in K18-hACE2 mice. K18-hACE2 mice (n=35) were inoculated intranasally with 1 x 10^6^ plaque forming units (PFU). Body weight (A), clinical signs (B), temperature (C), and survival (D) were monitored daily in sham/PBS animals (black, up to 7 dpi) and in infected animals (male, red; female, blue; up to 14 dpi). Animals meeting euthanasia criteria were counted dead the following day. Viral loads (genome copy numbers/mg) or infectious virus particles (PFU/mg of tissue) were monitored in the lung and brain (E-F). RNA copies were also examined in the serum (genome copies/mL) either directly on serum (G) or via a re-infectivity assay (H) using Vero E6 cells. The limit of detection is shown with a dashed line. n=3 (sham/PBS, n=19 (male), n=16 (female). One-way or two-way ANOVA. \**p*≤0.05, \*\**p*≤0.01, \*\*\**p*≤0.001, \*\*\*\**p*≤0.0001. For A-D, blue and green asterisks compare sham group vs. male group, and sham group vs. female group respectively. NA, non-applicable statistical test due extensive animal death at 6 dpi and limited n at 7 dpi.

Peak of lethality was associated with significant increases in viral loads (viral RNA and infectious virus particles) in the brain of the K18-hACE2 mice (Fig. 1E,F), as previously reported (19, 38, 41). In the lung, viral RNA copies were detectable at the earliest experimental timepoint (2 dpi) and remained stable over time, consistently within the value range reported in previous studies (30). While viral RNA remained high, viral titers however gradually declined over time, with no infectious virus recovered by 7 dpi (Fig. 1E-F). Although the absence of recoverable infectious virus in the lungs at 7 dpi was unexpected, this sharp decline mirrors published work that illustrated a 100-fold decline in PFUs at 7 dpi compared to 4 dpi (30). In contrast, viral RNA and infectious particles in the brain were low to undetectable at 4 dpi, but dramatically increased at 7 dpi (Fig 1E-F) representing the highest mean viral RNA load and infectious virus particles during the study. A small amount of viral RNA was detected in the serum (Fig.1G); however, incubation of SARS-CoV-2 permissive cell lines with serum samples did not result in any detectable productive infection *in vitro*, confirming an absence of viremia in intranasally-inoculated K18-hACE2 mice (Fig. 1H). Altogether, our data illustrate that lethality was associated with increasing viral RNA loads and infectious virus particles in the brain, which were simultaneously declining in the lung.

### SARS-CoV-2 results in transient mild infection in the nasal cavity of K18-hACE2 mice

We next performed detailed histologic analysis of various tissues to uncover the morphologic correlates of lethality in K18-hACE2 mice. For this, we first focused on the spatial and temporal dynamics of SARS-CoV-2 infection in the upper respiratory tract and analyzed the anterior/rostral nasal cavity (Fig. 2A-F) and olfactory neuroepithelium (Fig. 2G-L) for disease-associated lesions. To do so, we performed a thorough sequential histologic analysis combined with immunohistochemistry (IHC) and RNAscope® *in situ* hybridization (42) to determine the cellular localization and abundance of SARS-CoV-2 protein and RNA, using an anti-spike monoclonal antibody and an S-specific RNA probe, respectively.

**Fig. 2.**
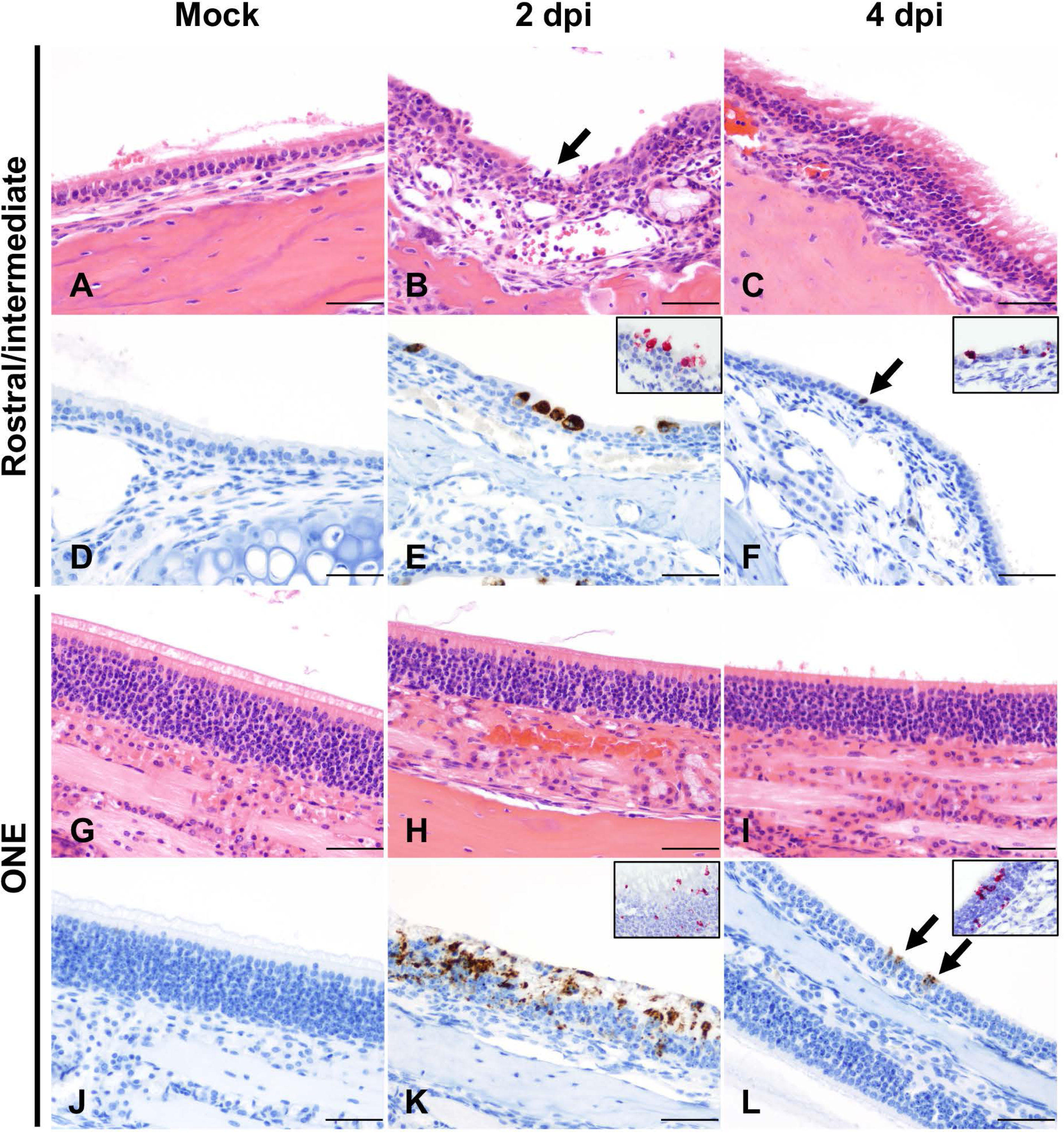
Temporal analysis of SARS-CoV-2 infection in the nasal cavity of K18-hACE2 mice. Histological changes, and viral protein (brown) and RNA (red) distribution and abundance were assessed in non-infected (mock: A, D, G, J) and infected mice at 2 (B, E, H ,K) and 4 (C, F, I, L) days following intranasal inoculation. At 2 dpi, suppurative rhinitis in the rostral and intermediate turbinates (B, arrow) correlated with abundant intraepithelial SARS-CoV-2 protein (E) and RNA (E, inset). Abundant viral protein and RNA were detected in the olfactory neuroepithelium (ONE, K and inset) in the absence of histologic lesions (H). At 4 dpi, only sporadically infected cells were noted in the epithelium lining the nasal turbinates and ONE (F and L, arrow and insets) in the absence of histologic lesions (C and I). Mock-infected are depicted in A, D, G and J. H&E and Fast Red (viral RNA), 200X total magnification. Bar = 100 μm.

At 2 dpi, the anterior/rostral nasal cavity was characterized by mild, multifocal neutrophilic inflammation (rhinitis) with segmental degeneration and necrosis of transitional and respiratory epithelium (Fig. 2B), which colocalized with intracytoplasmic SARS-CoV-2 protein and RNA (Fig. 2E). Adjacent nasal passages were partially filled with small amounts of cellular debris, degenerate neutrophils, and small numbers of erythrocytes. The lamina propria underlying affected areas was infiltrated by low to mild numbers of neutrophils and fewer lymphocytes (Fig. 2B). At 4 dpi, epithelial degeneration and necrosis in the rostral and intermediate turbinates was no longer observed, replaced by mild residual lymphocytic rhinitis and rare neutrophils within the lamina propria (Fig. 2C), and absence of exudate within nasal passages. SARS-CoV-2 protein and RNA were less commonly observed and restricted to rare positive cells in the respiratory epithelium (Fig. 2F and Table 1 and Table S1). By 7 dpi, the anterior/rostral nasal cavity was histologically within normal limits and no SARS-CoV-2 protein or RNA were detectable (Fig. 2C, F and Table 1 and Table S1).

**Table 1.** SARS-CoV-2 viral protein abundance in tissues derived from SARS-CoV-2-infected K18-hACE2 mice. Median scores are represented along with ranges between brackets when applicable.

| DPI | AT1/AT2 | Bronchioles | Rostral turbinates | Intermediate turbinates | ONE | Olf. bulb | Brain | Spinal cord (CT) | Spinal cord (LS) | GI* | Kidneys |
| --- | --- | --- | --- | --- | --- | --- | --- | --- | --- | --- | --- |
| Mock | 0 | 0 | 0 | 0 | 0 | 0 | 0 | 0 | 0 | 0 | 0 |
| 2 | 2 (1-2) | 0 | 1 (0-2) | 2 (1-2) | 1 (1-2) | 0 | 0 | 0 | 0 | 0 | 0 |
| 4 | 2 (1-3) | 0 | 0 (0-1) | 0 (0-1) | 1 (0-1) | 0 (0-1) | 0 (0-1) | 0 | 0 | 0 | 0 |
| 6-8 | 2 (1-3) | 0 | 0 | 0 | 1 (0-1) | 1 (0-2) | 3 (0-3) | 1 (0-2) | 0 (0-1) | 0 | 0 |
| 14 | 0 | 0 | 0 | 0 | 0 | 0 | 0 | 0 | 0 | 0 | 0 |

0, no SARS-CoV-2 protein observed; 1, 0 to 5% of cells within a high magnification (400X) field are positive for viral protein; 2, 5 to 25% of cells within a high magnification (400X) field are positive for viral protein; 3, >25 to <50% of cells within a high magnification (400X) field are positive for viral protein. NA, not available. AT1, alveolar type 1 pneumocytes; AT2, alveolar type 2 pneumocytes; ONE, olfactory neuroepithelium; CT, cervicothoracic segment; LS, lumbosacral segment; GI, gastrointestinal tract.
\*Sections examined included stomach, small intestine (duodenum, jejunum, and ileum) and large intestine (cecum and colon).

The posterior nasal cavity, olfactory neuroepithelium (ONE) (Fig.2G-L), displayed mild segmental degeneration and necrosis at 2 dpi, which colocalized with abundant SARS-CoV-2 protein and RNA (Fig. 2K and Table 1 and Table S1). By 4 dpi, histopathologic lesions in the ONE had resolved, but rare SARS-CoV-2 protein and RNA were observed both at 4 and 7 dpi (Fig. 2L and Table 1). No SARS-CoV-2 protein or RNA were detected in the ONE by 14 dpi (Table 1 and Table S1).

### SARS-CoV-2 induces moderate interstitial pneumonia in K18-hACE2 mice

In the lower respiratory tract, histologic alterations in the pulmonary parenchyma mainly involved the alveoli, interstitium and perivascular compartment (Fig. 3A-K). Overall, pathologic alterations in the lungs were characterized by moderate progressive lymphohistiocytic and neutrophilic interstitial pneumonia that peaked at 7 dpi (Fig. 3G, H). We quantitatively analyzed the total % of pneumonia using a machine learning classifier to differentiate normal vs. pneumonic lung tissue. Peak disease was confirmed to occur at 7 dpi, with a mean of ∼10% of total lung area affected, with one outlier of ∼40% of total lung area, suggesting that more severe disease is possible, albeit uncommon (Fig. 3K).

**Fig. 3.**
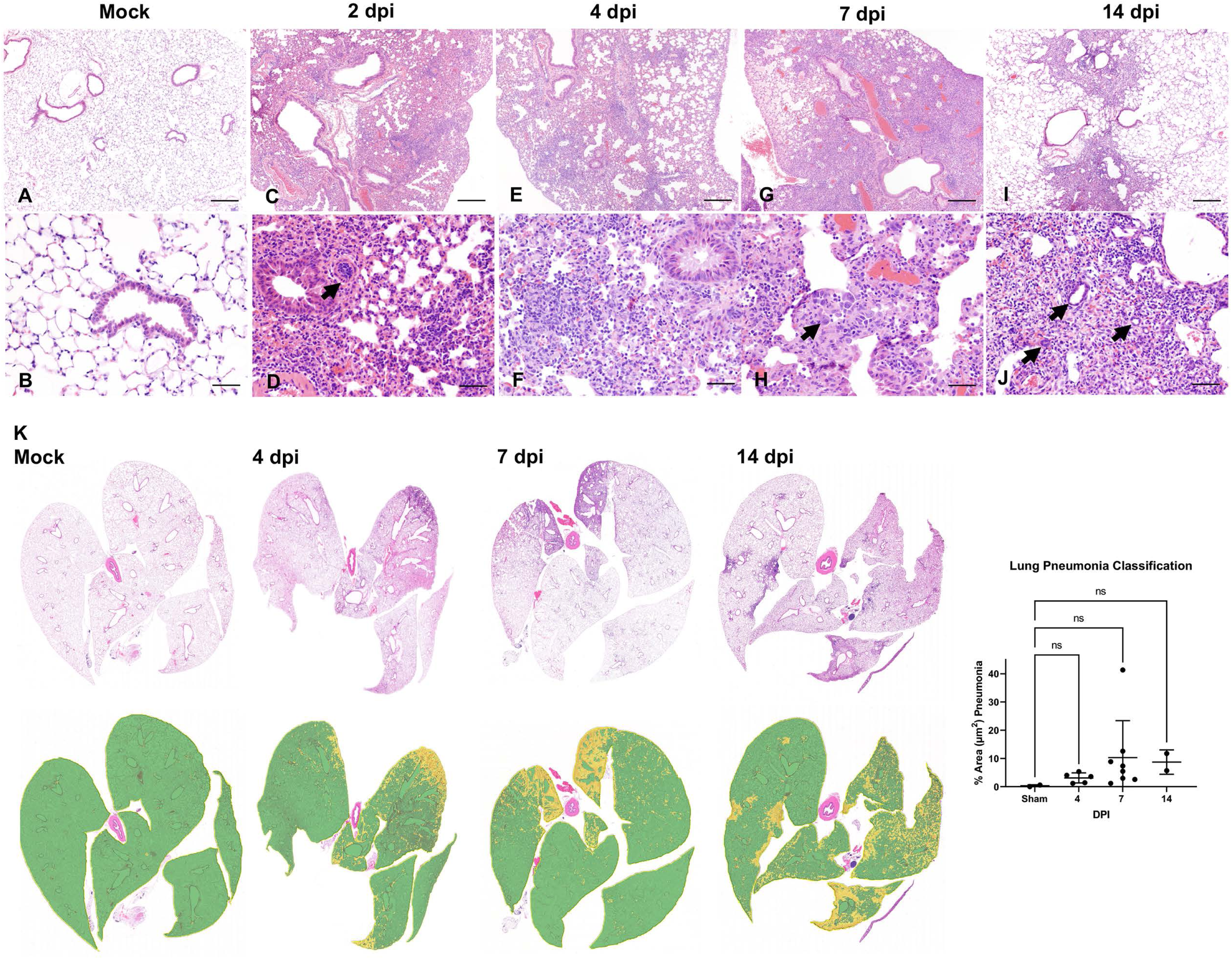
Temporal analysis of SARS-CoV-2 infection in the lungs of K18-hACE2 mice. Lung tissues from non-infected mice (Mock: A, B) and from infected mice at 2 (C, D), 4 (E, F), 7 (G, H) and 14 (I, J) days following intranasal inoculation were analyzed. Subgross histological images of the lungs and corresponding pneumonia classifiers for each timepoint are depicted in panel K (green = normal; yellow = pneumonia). Mild to moderate interstitial pneumonia was evident starting at 2 dpi with frequently reactive blood vessels (D, arrow). At 7 dpi, alveolar type 2 (AT2) cell hyperplasia was observed (H, arrows). Residual mild pneumonia was observed in the rare animals that survived to 14 dpi, with rare sporadic lymphoid aggregates (J-arrows). H&E, 50X (A, C, E, G, and I; bar = 500 μm), 200X (B, D, F, H and J; bar = 100 μm) and 1X (K) total magnification. One-way ANOVA; ns, non-significant.

Of note, data from our 2 dpi animals were excluded from tissue classification analysis as lungs were sub-optimally insufflated and the classier algorithm falsely labeled areas of atelectasis as pneumonia. Pneumonia was interpreted to be minimal at this timepoint and, thus, considered negligible.

At 2 dpi, minimal perivascular and peribronchiolar inflammation, consisting primarily of lymphocytes and histiocytes, and occasional perivascular edema were observed (Fig. 3C). Pulmonary vessels were frequently reactive and lined by a plump endothelium with marginating leukocytes (Fig. 3D). SARS-CoV-2 protein and RNA (Fig. 4A-J) were observed in proximity to areas of interstitial pneumonia and localized within the cytoplasm of alveolar type (AT) 1 (squamous epithelium) and fewer AT2 cells (cuboidal epithelium) (Fig. 4B, G).

**Fig. 4.**
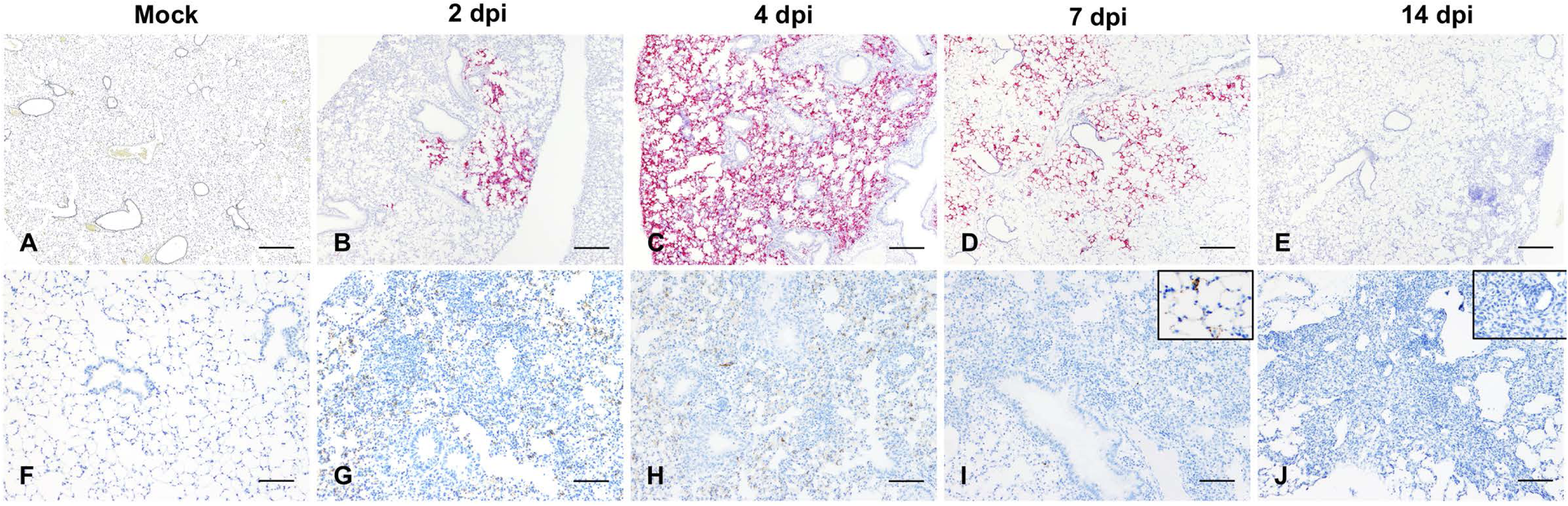
Temporal analysis of SARS-CoV-2 RNA and protein distribution in the lungs of K18-hACE2 mice. Presence of viral RNA (A-E) and Spike protein (F-J) was assessed by ISH and IHC in non-infected (mock: A, F) and infected lung tissues from K18-hACE2 mice at 2 (B, G), 4 (C, H), 7 (D, I) and 14 (E, J) days following intranasal inoculation. Peak viral RNA and protein occurred at 4 dpi (C,H), with an evident decline by 7 dpi (D, I). No viral RNA or protein were detected at 14 dpi (E, J). Fast Red, viral RNA (A-E) and DAB viral protein (F-J), 100X total magnification. Bar = 200 μm. Insets (I&J), 400x total magnification.

At 4 dpi, peak in viral protein and RNA abundance were observed (correlating with the highest viral titer and RNA load as determined by RT-qPCR and plaque assays) (Fig 4C, H) along with increasing lymphohistiocytic and neutrophilic infiltrate (Figs. 3E, F). SARS-CoV-2 cellular tropism did not differ from that described at 2 dpi, but SARS-CoV-2 protein and RNA were more abundant and routinely observed in histologically normal pneumocytes (Fig 4C, H).

At 7 dpi, lymphohistiocytic and neutrophilic interstitial pneumonia peaked in severity, which on average was moderate to regionally severe, affecting ∼10-40% of the parenchyma (Fig. 3G, H, K). Additional unique findings at 7dpi included, rare alveolar septal necrosis, mild proliferation of AT2 cells, and sporadic regional pulmonary edema (Fig. 3G, H). SARS-CoV-2 protein and RNA were occasionally still abundant in several animals, but predominated in histologically normal parenchyma, with minimal to rare detection in areas of prominent inflammation (Fig. 4D, I and Table 1, and Table S1). Taken together, this suggested progressive resolution of viral infection by the host consistent with declining viral loads and absence of culturable infectious virus at this timepoint (Fig. 1E, F).

In the two survivors euthanized at 14 dpi, persistent mild to moderate lymphohistiocytic interstitial pneumonia was observed, with formation of sporadic lymphoid aggregates and mild persistence of AT2 hyperplasia (Fig. 3I, J). SARS-CoV-2 protein or RNA were no longer detectable at 14 dpi (Fig 4E, J).

Of note, no evidence of SARS-CoV-2 infection was observed in bronchiolar epithelium and pulmonary vasculature at any time during the study (Figs. 3 and 4, and Table 1, and Table S1). Similarly, hyaline membranes, vascular thrombosis, and syncytial cells were not observed at any time point across all animals, which contrasts with serve disease described in human autopsies (43) and non-human primate studies (44, 45). In one animal (7 dpi), there was localized flooding of bronchioles by degenerate neutrophils and cellular debris mixed with birefringent foreign material consistent with aspiration pneumonia, a rare complication previously reported in K18 hACE2 mice infected with SARS-CoV-1 that was ultimately attributed to pharyngeal and laryngeal dysfunction secondary to central nervous system (CNS) disease (36).

Altogether, our data displays evidence of a significant but moderate lymphohistiocytic interstitial pneumonia in SARS-CoV-2 infected K18-hACE2 mice. Histopathological features contrast with those observed in severe cases of COVID-19 in humans and suggest that the lethality observed in this model is in part independent of virally induced lung injury and resultant pneumonia.

### Pulmonary SARS-CoV-2 replication occurs exclusively in AT1 and AT2 cells

Subsequently, we aimed to further investigate SARS-CoV-2 tropism in the lower respiratory tract of K18-hACE2 mice. We first performed qualitative multiplex IHC to probe the localization of SARS-CoV-2 protein in AT1 cells (cell maker: receptor for advanced glycation end-products [RAGE]), AT2 cells (cell marker: surfactant protein C [SPC]), and endothelial cells (cell marker: CD31). SARS-CoV-2 protein was restricted within RAGE+ AT1 and SPC+ AT2 pneumocytes, but not with CD31+ endothelial cells (Fig. 5A-C). Transmission electron microscopy (TEM) corroborated our IHC and ISH data, where we observed viral protein and RNA exclusively within the cytoplasm of squamous and cuboidal pneumocytes. Double membrane-bound vesicles (DMVs) and virus particles were exclusively observed in cells containing abundant caveolae (AT1 cells) or lamellar bodies (AT2 cells) by transmission electron microscopy (Fig. 5D-F). No viral particles or replication intermediates were observed in bronchiolar epithelial cells (Fig. 5G, H) or vascular endothelium. Of note, cubic membranes were a distinctive feature only observed in AT1 pneumocytes (Fig 5E).

**Fig. 5.**
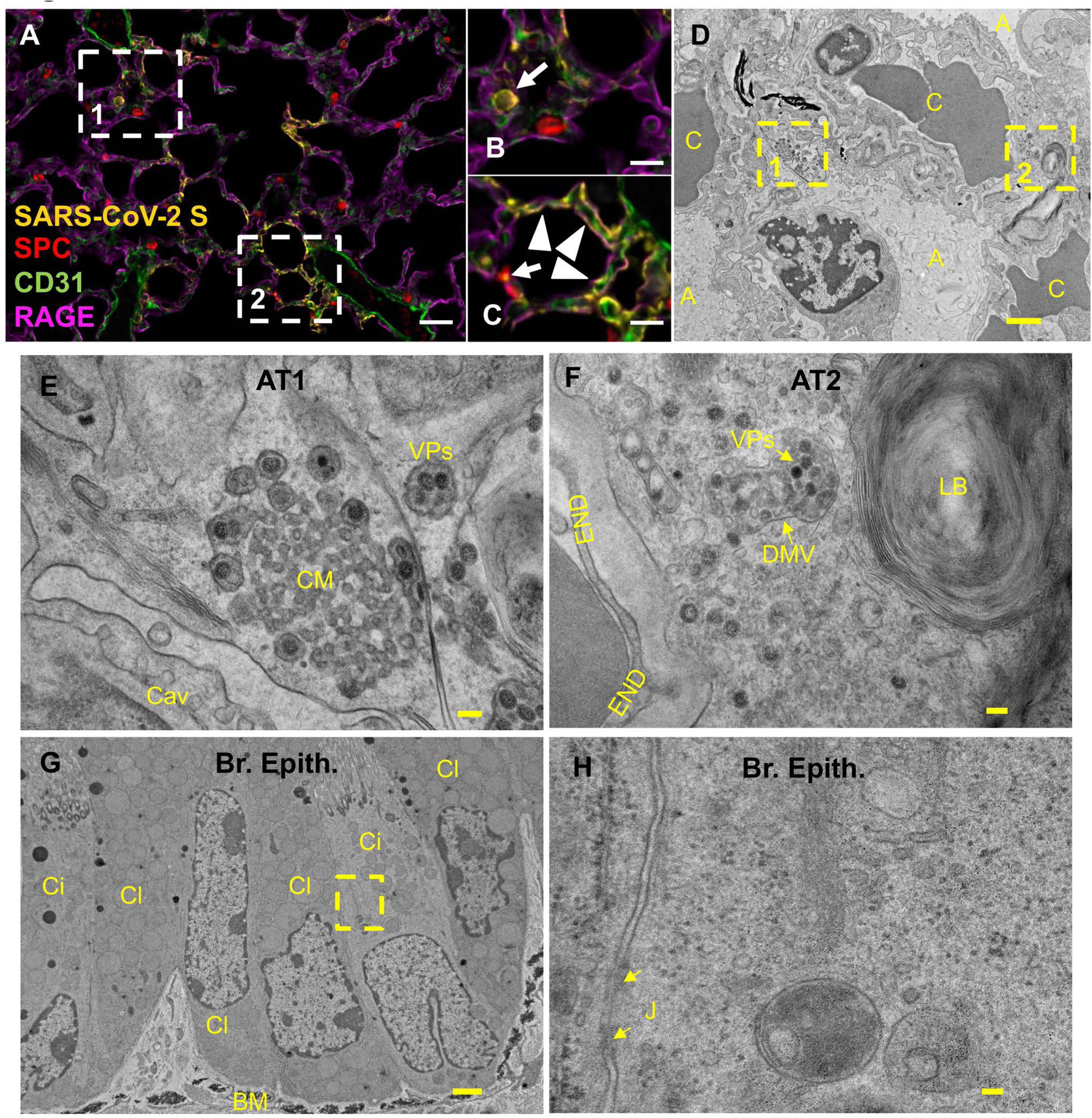
SARS-CoV-2 tropism following intranasal inoculation in K18-hACE2. (A-C) At 4 dpi, SARS-CoV-2 (yellow) showed tropism for RAGE^+^ alveolar type 1 (AT1, magenta) and scattered SPC^+^ alveolar type 2 (AT2, red) cells (B and C, arrowheads, and arrows, respectively) but not for CD31^+^ endothelial cells. B and C represent magnification of inset 1 and 2 from A, respectively. (D-F) 6 dpi, virus particles (VPs) were bound by double membrane vesicles (DMVs) in AT1 (E) and AT2 (F) cells. AT1 contained abundant caveolae. Another unique feature observed in AT1 cells was the presence of cubic membranes (CM). AT2 pneumocytes were characterized by presence of lamellar bodies. E and F represent magnification of inset 1 and 2 from D, respectively. (G, H) Viral particles were not identified in ciliated or non-ciliated club bronchiolar epithelium. H represents an inset magnification of G. Multiplex fluorescent IHC, 100x (A; bar= 100 μm) and 200x (B,C; bar = 50 μm) total magnification. TEM, bar = 2 μm (B), 100 nm (B1, B2 and C1), and 3 μm (C). A, alveolar lumen; BM, basement membrane; C, capillary; Cav, caveolae; Ci, ciliated epithelium; Cl, club epithelium; CM, cubic membranes; DMVs, double-membrane vesicles; END, endothelium; J; cell-cell junction; VPs, viral particles.

### Effective control of SARS-CoV-2 infection in the lower respiratory tract is associated with recruitment of macrophage and to a lesser degree cytotoxic T cells

Next, we quantitatively characterized the cell density of inflammatory cells (cells/μm^2^) targeting macrophages (Iba1), cytotoxic T cells (CD8), B cells (CD19) and total area immunoreactivity (% area μm^2^) of viral protein (Spike) in the lungs of SARS-CoV-2 infected K18-hACE2 mice (Fig. 6A-H). SARS-CoV-2 S immunoreactivity peaked between 4-7 dpi (Fig. 6A), supporting a positive correlation between viral infection and the progressive inflammatory cell infiltrate, but was not statistically significant across groups. We attribute this finding to our low sample size for quantitative whole slide analysis, especially at 2 and 4 dpi, and individual animal variability likely represented by the inherent heterogeneity of viral pneumonia. Iba1+ macrophages represented the predominant inflammatory infiltrate across all time points with a temporal increase peaking at 7 dpi (p=0.0044 compared to sham inoculated mice, Fig. 6B, G). Cytotoxic T cells were the second most abundant inflammatory infiltrate quantified, which also displayed a temporal increase peaking around 4-7 dpi (Fig. 6C, F-G); however, these cells were present at a ∼10-fold reduced frequency compared to macrophages and statistical significance was not observed across timepoints, suggesting an early and plateaued response of this inflammatory population. B cells were elevated by 7 dpi but reached peak cell density at 14 dpi (Fig. 6D, H), the only time point where discrete lymphoid aggregates were observed histologically (p≤0.0001 compared to sham inoculated mice). Altogether, our data suggest that a strong and persistent myeloid infiltrate and, to a lesser degree, cytotoxic T cells are important contributors to the decline of viral load that occurs in the lungs between 4-7 dpi, with B cells potentially being involved if animals survive the acute stage of disease. Of note, minimal inflammatory cells were observed in sham-inoculated K18-hACE2 mice supporting that the two survivors were de-facto infected with SARS-CoV-2 (Fig. 6E), which was further supported by the presence of neutralizing antibodies in their serum as compared to naïve mice (Fig. 6I).

**Fig. 6.**
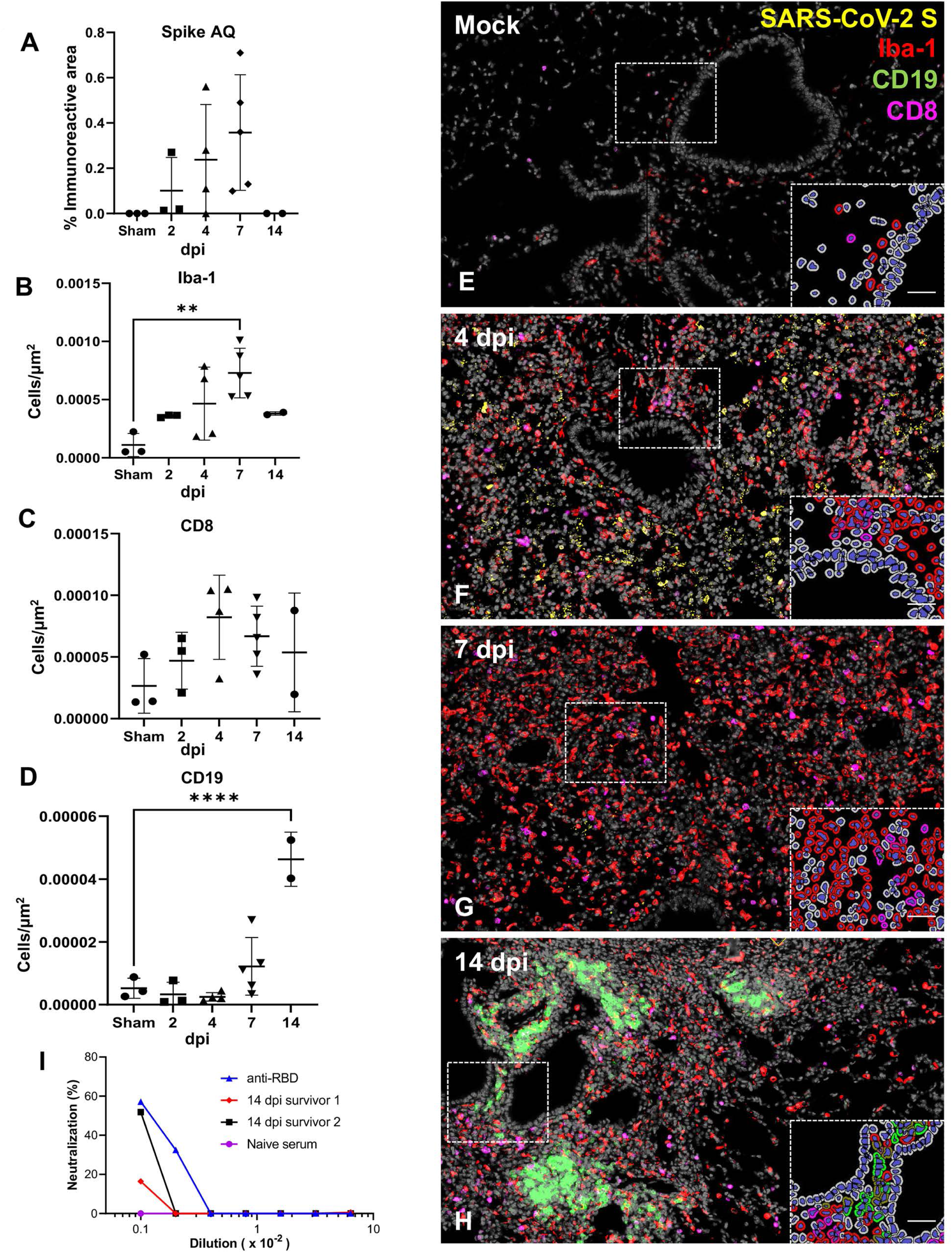
Temporal immunoprofiling of the pulmonary host inflammatory response to SARS-CoV-2. (A-H) Quantification and 4-plex fluorescent IHC targeting SARS-CoV-2 Spike (A, E-H), and macrophage Iba-1+ (B, E-H), CD8+ (C, E-H) and CD19+ cell (D, E-H) infiltration in the lung of sham/PBS mice and in inoculated mice (2, 4, 7 and 14 dpi). In inoculated mice, SARS-CoV-2 Spike peaked between 4-7 dpi (A, F-G). Iba-1+ macrophages (red) increased significantly peaking at 7 dpi (B, G), along with a lower infiltration of CD8+ T lymphocytes-magenta that peaked between 4-7 dpi (C,F-G) while Sham/PBS mice had low residual inflammatory cells (E). CD19+ B cells arranged in clusters were only evident in the two survivors euthanized at 14 dpi (D, H). Insets depict immune cell phenotyping outputs that were applied across the entire whole slide. Multiplex fluorescent IHC (E-H): 100X and 400X (insets) total magnification, bar=50μm. I. Neutralizing activity of serum isolated from a naïve/non-infected K18-hACE2 (purple) and from the two 14 dpi survivors at 14 dpi (survivor 1 and 2, red and black respectively). An anti-SARS-CoV-2 Spike RBD antibody (anti-RBD, blue) was used as a positive control. Serum was serially diluted by 2-fold. One-way ANOVA. \*\**p*≤0.01.

### SARS-CoV-2 exhibits extensive neuroinvasion with resultant neuronal degeneration and necrosis in K18-hACE2 mice

Pursuing our hypothesis that the lethality of the K18-hACE2 mice is associated with neuroinvasion, we analyzed sagittal sections of the whole head to characterize progression of histologic lesions and distribution of viral protein and RNA at different timepoints post-infection (2, 4, 6-7 and 14 dpi). Histologic alterations in the brain were severe and widespread by 7 dpi with involvement of the olfactory bulb and of the cerebrum (Fig.7), as well as of the cerebral cortex (most predominantly somatosensory and somatomotor areas), hippocampus (mainly CA1 region), midbrain (thalamus and hypothalamus), brainstem, and the dentate nucleus. Histologic findings included moderate to marked neuronal spongiosis, multifocal shrunken, angular, hypereosinophilic and pyknotic neuronal bodies with loss of Nissl substance/chromatolysis (neuronal degeneration and necrosis, Fig. 7J) and occasionally delimited by multiple glial cells (satellitosis). In the olfactory bulb, delicate lymphocytic perivascular cuffing (Fig. 7K) and a general increase in the number of reactive glial cells (gliosis) were evident in the neuroparenchyma neighboring areas of neuronal degeneration and necrosis. Notably, the cerebellum (cortical layers and associated white matter of the cerebellar folia) was spared of histologic changes (data not shown).

**Fig. 7.**
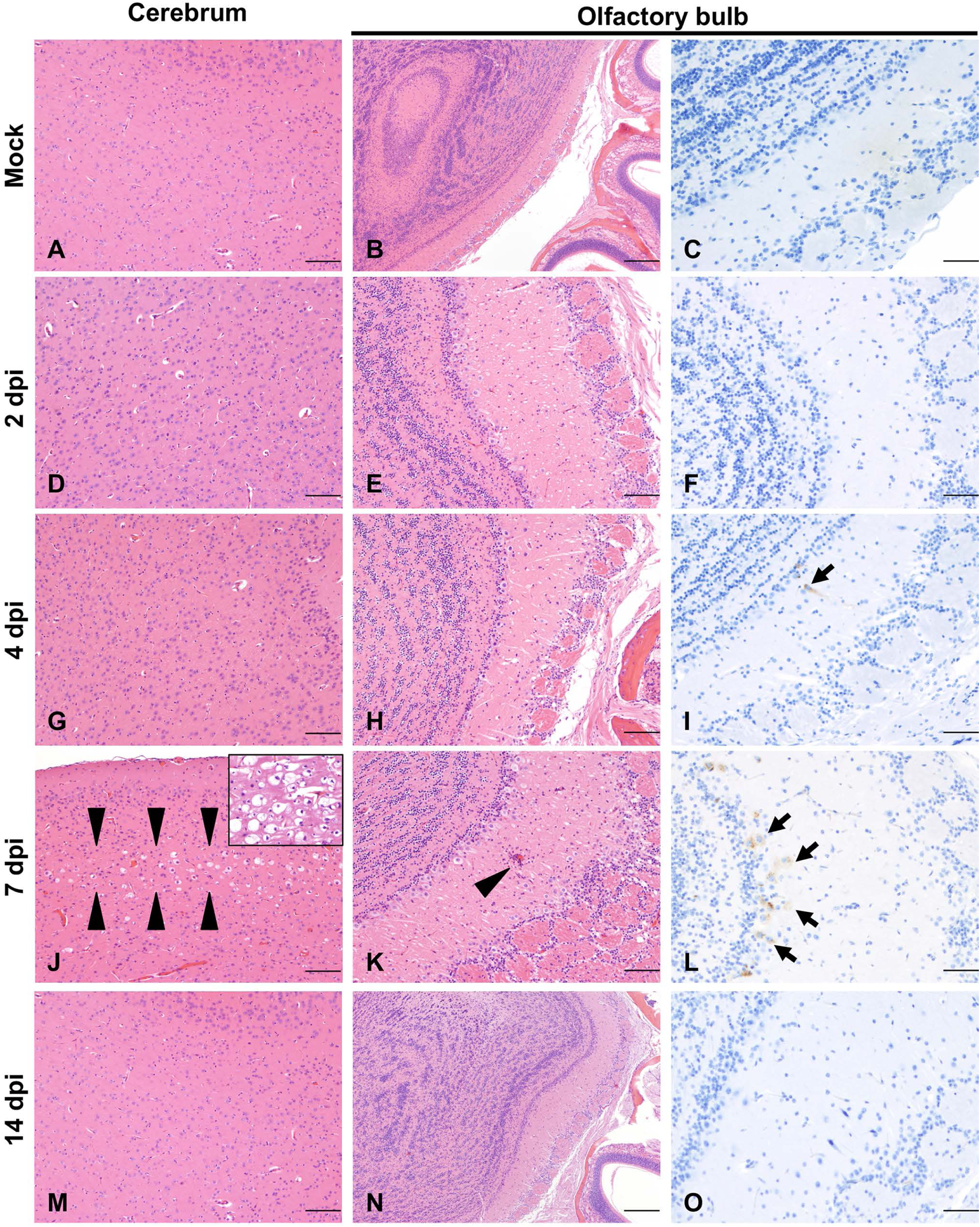
Temporal neuronal damage in K18-hACE2 mice following intranasal inoculation with SARS-CoV-2. Histologic changes in the cerebrum (A, D, G, J, M) or olfactory bulb (B, E, H, K, N) in non-infected or infected K18-hACE2 mice. SARS-CoV-2 Spike protein was also probed in the olfactory bulb of non-infected and infected K18-hACE2 mice by IHC (C, F, I, L, O). SARS-CoV-2 protein (brown) was evident as early as 4 dpi (I, arrow), but no histologic changes were noted until 7 dpi (J-K). At that time point, mild (J, arrowheads) to marked (J, inset) spongiosis with neuronal degeneration and necrosis involving multiple areas within the cerebral cortex and elsewhere were observed. Similar changes were evident in the olfactory bulb, with occasional perivascular cuffs/gliosis (K, arrowhead) and abundant viral protein (L, arrows). No histologic alterations or viral protein were detected in survivor mice euthanized at 14 dpi (M-O). H&E and DAB (viral protein), 100X (A, B, D, E, G, H, J, K, M, N; bar = 200 μm) and 200X (C, F, I, L, O; bar = 100 μm) total magnification.

Neuronal morphologic changes directly correlated with abundant neuronal immunoreactivity for SARS-CoV-2 S protein and viral RNA, which was observed exclusively within neuronal cell bodies and processes (Figs. 7C, F, I, L, O, and 8A). SARS-CoV-2 protein and RNA had a widespread distribution throughout the brain in roughly 85% (11/13) of infected K18-hACE2 mice at 7 dpi, including neuronal bodies within the cerebral cortex, CA1, CA2 and CA3 regions of the hippocampus, anterior olfactory nucleus, caudoputamen, nucleus accumbens, thalamic nuclei including hypothalamus, midbrain, pons and medulla oblongata nuclei (Fig. 8A). Few vestibulocochlear nerve fascicles showed immunoreactivity for viral protein; while no viral S protein or RNA was detected in areas spared of histological changes including the cerebellar cortex and white matter, optic nerve and retina, and the spiral ganglion of the inner ear (albeit the eye and inner ear were not present in most sections examined). SARS-CoV-2 S protein and RNA preceded histological findings with rare detection as early as 4 dpi in mitral and inner nuclear neurons of the olfactory bulb, as well as small clusters of neurons within the anterior olfactory nucleus and orbital area of the cerebral cortex (Figs. 7C, F, I, L, O, and 8A).

**Fig. 8.**
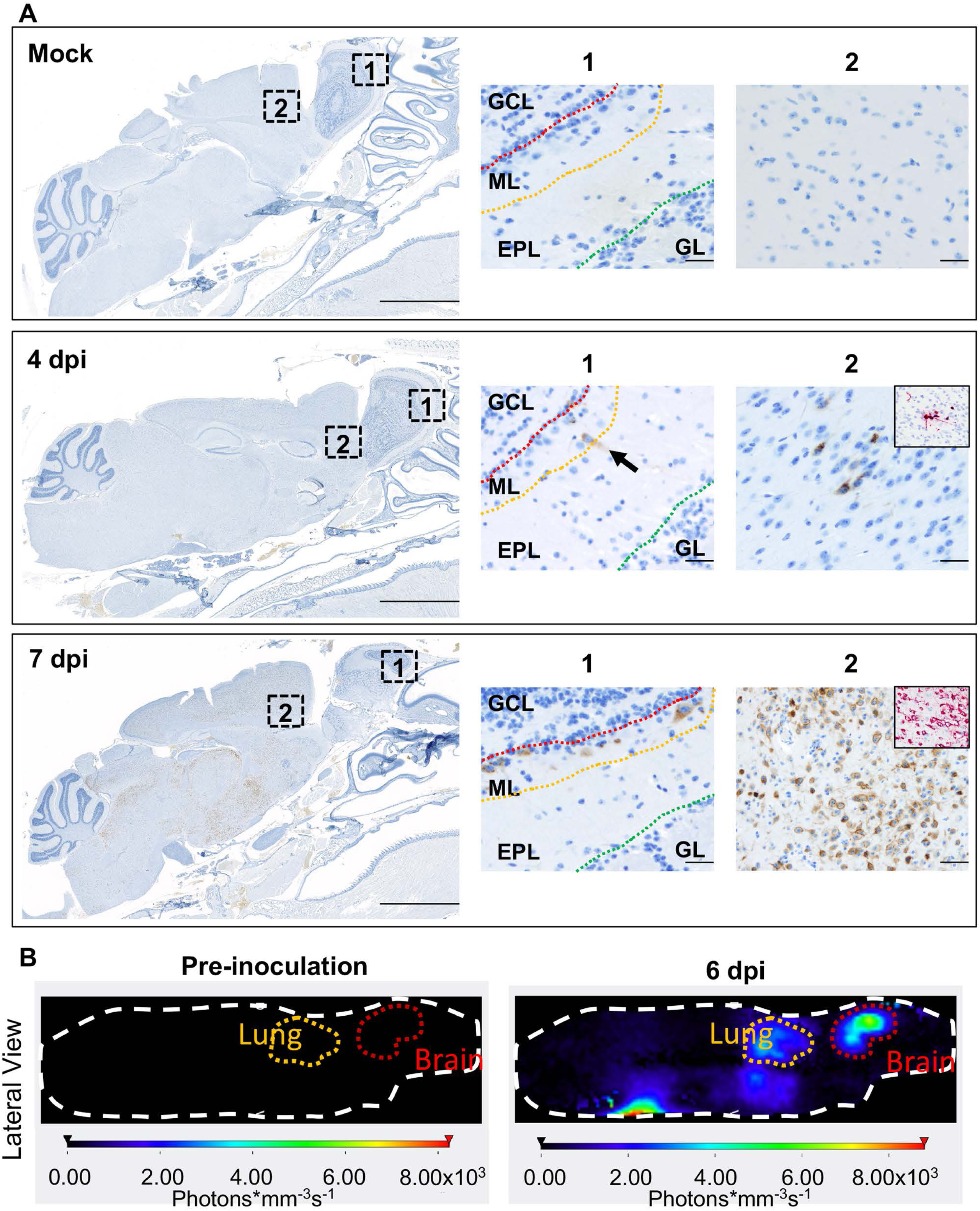
Invasion of SARS-CoV-2 into the central nervous system. (A)Sagittal sections of the head of non-infected (mock, top panel) and infected (4 and 7 dpi, middle and bottom panel respectively) were analyzed for viral protein and RNA distribution. At 4 dpi (middle panel), SARS-CoV-2 infected neurons within the mitral layer of the olfactory bulb (1, arrow) as well as small clusters of neuronal bodies within the cerebral cortex (2, SARS-CoV-2 RNA in inset). At 7 dpi (bottom panel), SARS-CoV-2 protein was widespread along the mitral layer of the olfactory bulb (1) and throughout the central nervous system (2, SARS-CoV-2 RNA in inset) with exception of the cerebellum. EPL, external plexiform layer; GCL, granular cell layer; GL, glomerular layer; ML, mitral layer. DAB (viral protein) and Fast Red (viral RNA). 7.5X (bar = 2.5 mm) and 200X (bar = 100 μm) total magnification. On the right of each panel, pictures labelled 1 and 2 are 266X total magnification insets represented by the hashed squares labeled in the lower (7.5X) magnification images. (B) Representative three-dimensional profile view (right side) of a K18-hACE2 mouse following inoculation with a rSARS-CoV-2 NL virus (10^6^ PFU). NanoLuc bioluminescent signal was detected and quantified at 6 dpi following fluorofurimazine injection (Sub-cutaneous) using the InVivoPLOT (InVivoAx) system and an IVIS Spectrum (PerkinElmer) optical imaging instrument. Location of the lungs and brain are indicated.

Using a NanoLuc expressing recombinant SARS-CoV-2 virus (rSARS-CoV-2 NL), we observed significant detection of bioluminescence in the brain of a representative K18-hACE2 mice at day 6 post-infection, which was associated with lower bioluminescence signal in the lungs at the same time point, consistent with our previous findings (Fig. 8B) and other reports (46). To better characterize the gliosis that was observed histologically in animals with abundant neuronal degeneration and necrosis we quantitatively characterized the total area immunoreactivity (% area μm^2^) of microglia (Iba-1), astrocytes (GFAP), and SARS-CoV-2 (S) in sagittal sections of whole brain (Fig. 9A-C). Total % area immunoreactivity for astrocytes (GFAP) and microglia (Iba1) dramatically increased at 7 dpi compared to sham inoculated negative controls (GFAP, p=0.0101; Iba1, p=0.0327), paralleling peak expression of SARS-CoV-2 S, which was also significantly increased compared to Sham inoculated negative controls (p=0.0351; Fig. 9A-C). Notably, the morphology of microglial, which had delicate cytoplasmic processes in sham inoculated (Fig. 9B), 2 and 4 dpi mice, was replaced by broad shortened processes at 7 dpi (Fig. 9C). Morphological differences in astrocyte processes were more subtle, but still possessed broader and a more extensive branching pattern compared to 2dpi, 4dpi, and sham inoculated negative controls (Fig. 9B, C). To confirm exclusive neuronal tropism, we performed TEM on an animal euthanized at 6 dpi that exhibited neurologic signs of disease in the form of tremors. Viral assembly was observed exclusively in neurons, with no detection in glial cells. The prominent histologic phenotype of neuronal spongiosis was characterized by profound accumulation of double membrane vesicles (DMVs) and virus particles, with loss of Nissl substance (degeneration) or nuclear pyknosis, karyolysis, and global electron dense transformation of the cytoplasm (necrosis) (Fig. 9D, E). Although viral assembly and/or particles were not observed in microglia or astrocytes, quantitative immunohistochemical analysis supports reactive microgliosis and astrogliosis that is temporally linked with peak neuroinvasion, suggesting that activation of these cells contributes to neuronal injury either through direct neurotoxic and/or loss of normal homeostatic neurotrophic mechanisms.

**Fig. 9.**
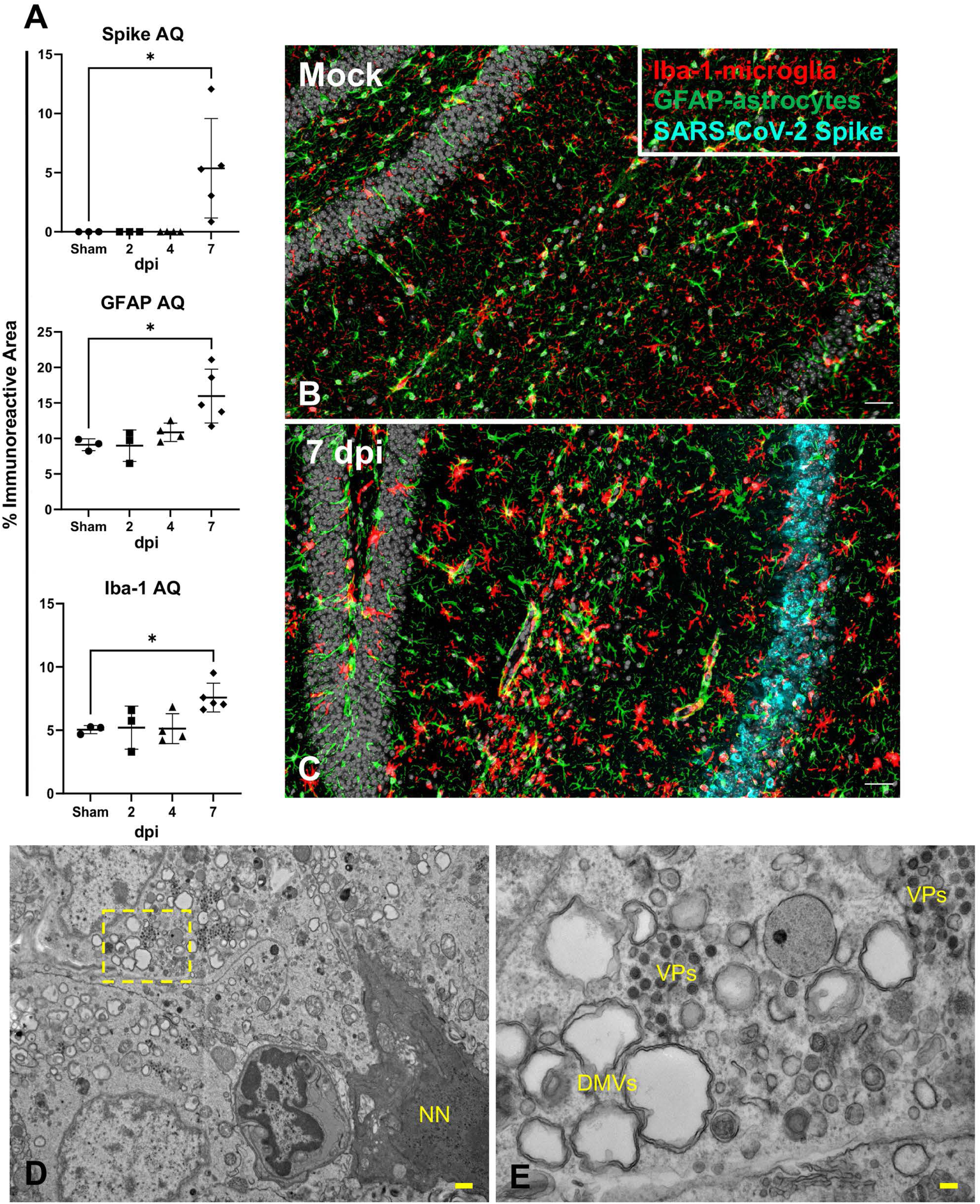
SARS-CoV-2 replication and assembly of virus particles in hippocampal neurons and glial response. (A-C) Quantification (A) of 3-plex fluorescent IHC (B,C) targeting SARS-CoV-2 Spike, astrocyte (GFAP) and Iba-1+ microglial infiltration in the brain of sham/PBS mice and in inoculated mice (2, 4, 7 and 14 dpi). The amount of viral protein rapidly and markedly increases by 7 dpi, along with an intense astrocytic and microglial response. (D-E) Ultrastructural examination of infected neighboring hippocampal neurons illustrated cytoplasmic swelling by numerous double membrane bound vesicles (DMVs) and free virus particles. Karyolysis and global electron dense transformation of the cytoplasmic compartment was also observed indicative of neuronal necrosis (NN). E is an inset magnification from D. Multiplex IHC, 200X total magnification, bar = 100 μm. TEM, bar = 100 nm (D) or 50nm (E). One-way ANOVA; \**p*≤0.05.

Considering the severe bladder distention noted at necropsy and proprioceptive deficits observed clinically, we examined the cervicothoracic and lumbosacral segments of the spinal cord. In 9/11 animals that died or were euthanized due to terminal disease, similar histologic findings were observed as those described in the brain, albeit with milder gliosis and lymphocytic perivascular cuffing (Fig. 10A). We also observed mild-to-moderate detection of viral protein in the spinal cord that predominated within neurons of the cervicothoracic segments (Fig. 10B, and Table 1 and Table S1). Finally, Luxol Fast Blue was utilized to visualize the integrity of myelin following SARS-CoV-2 invasion in the brain and spinal cord at 7 dpi, with no evidence of demyelination noted (Fig. 10C).

**Fig. 10.**
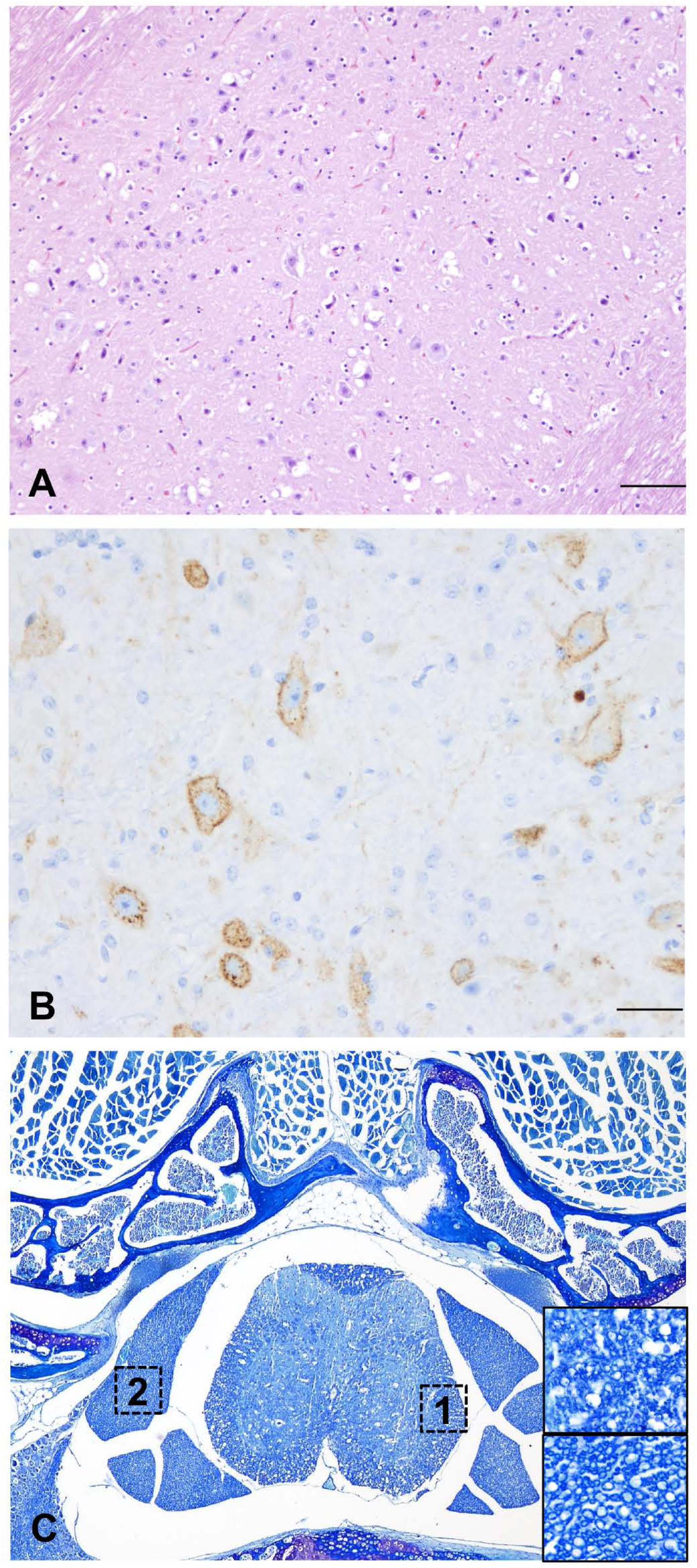
Histological and immunohistochemical findings in the cervicothoracic spinal cord of SARS-CoV-2-infected K18-hACE2 mice at 7 dpi. (A) At 7 dpi, multifocal neuronal bodies within the grey matter are shrunken, angular and hyperchromatic (neuronal degeneration and necrosis), and the neuroparenchyma has multiple clear spaces filled with small amounts of debris (spongiosis) with a slight increase in the number of glial cells (gliosis). H&E, 100X total magnification. (B) Abundant SARS-CoV-2 spike protein localized within the perikaryon and processes of neurons within the spinal cord. DAB, 200X total magnification. Bar = 100 μm. (C) No myelin loss was noted in the white matter. Luxol Fast Blue, 200X total magnification. Bar = 100 μm.

Taken together, our data illustrate that SARS-CoV-2 infection of K18-hACE2 results in severe neuronal invasion of the CNS, via transport to the olfactory bulb originating from axonal processes traversing the ONE. Alternative and concurrent routes such as retrograde transport of other cranial nerves as well as parasympathetic and sympathetic sensory nerves cannot be ruled out, especially acknowledging the near diffuse distribution of virus in the brain at terminal stages of disease, with the exception of the cerebellum. Viral neuroinvasion resulted in extensive neuronal cytopathic effect in infected cells that ultimately resulted in cell death, comprising not only the brain but also the spinal cord. Further research is warranted to characterize the role of uninfected but reactive microglia and astrocytes in SARS-CoV-2 neuronal injury using a multidimensional approach including molecular and functional testing.

### ACE2 expression and distribution does not fully reflect SARS-CoV-2 tissue tropism in K18-hACE2 mice

To further explore the mechanism driving lethal SARS-CoV-2 infection in K18-hACE2 mice, we first investigated the tissue and cellular distribution of the ACE2 receptor in both C57BL/6J and K18-hACE2 mice by IHC (Fig. 11A-L) using a cross-reactive anti-ACE2 antibody (cross-reactive to hACE2 and mACE2) (Table S2). In the lower respiratory tract (lungs), ACE2 was ubiquitously expressed along the apical membrane of bronchiolar epithelium and, less commonly, in rare and scattered AT2 pneumocytes (Fig. 11A-C). No ACE2 expression was found in AT1 pneumocytes. No evident differences in the distribution and abundance of ACE2 expression were identified between uninfected C57BL/6J, sham-inoculated K18-hACE2, and terminal (7 dpi) K18-hACE2 mice inoculated with SARS-CoV-2.

**Fig. 11.**
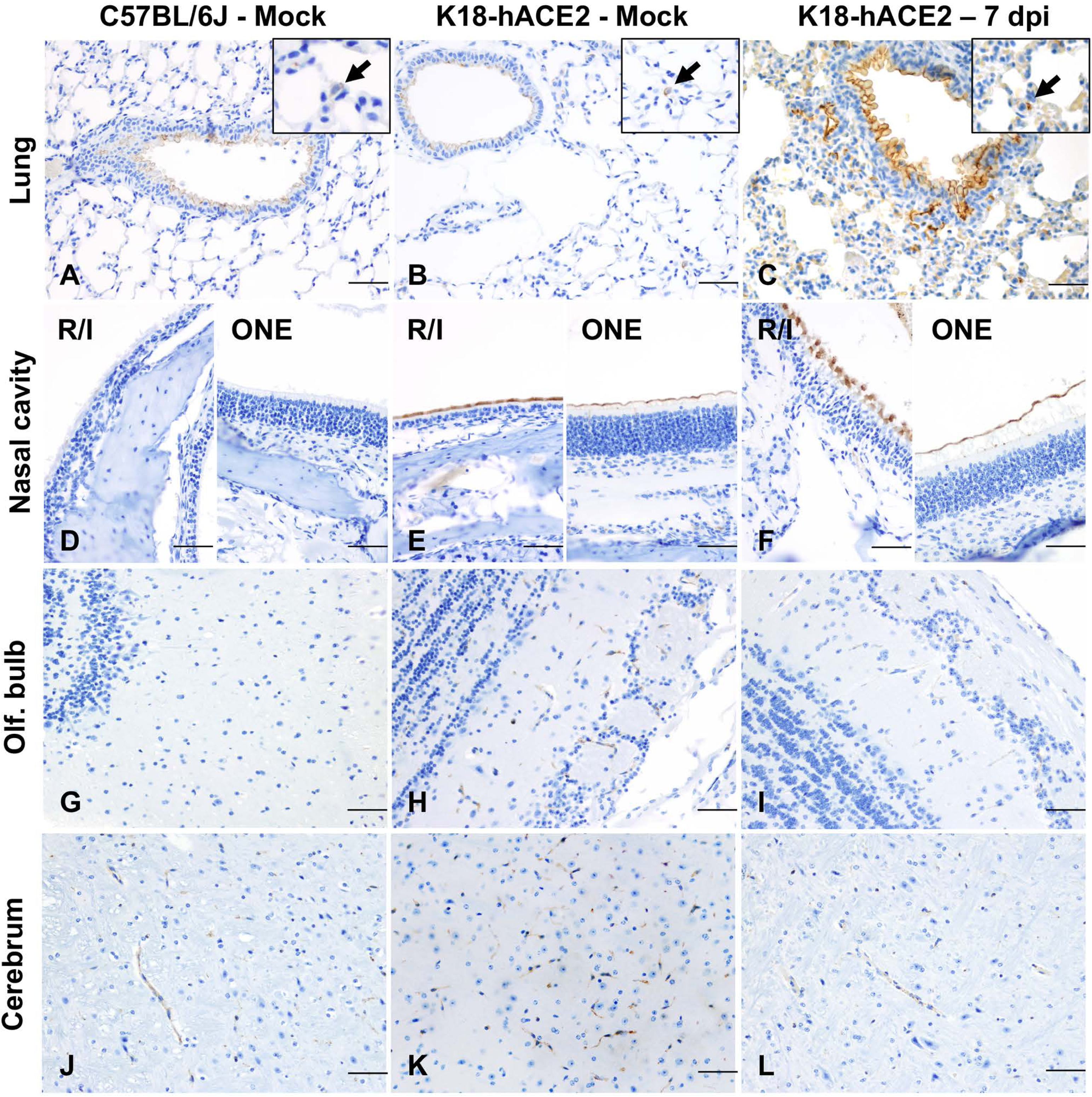
Distribution of ACE2 in lungs, nasal cavity, brain, and olfactory bulb of wild-type C57BL/6J and transgenic K18-hACE2 mice. Lung (A-C), nasal (rostral/intermediate turbinates [R/I]) and olfactory epithelium (ONE) (D-F), olfactory bulb (G-I) and brain (J-L) from non-infected C57BL/6J, and from non-infected and infected (7 dpi). K18-hACE2 mice were analyzed via immunohistochemistry using a cross-reactive anti-ACE2 antibody. In the lungs (A-C), ACE2 expression (brown) was mostly restricted to the apical membrane of bronchiolar epithelial cells with scattered positive AT2 cells (inset arrows). Nasal (rostral/intermediate turbinates [R/I]) and olfactory epithelium (ONE) were devoid of ACE2 in C57BL/6J mice (D) but expression was enhanced in K18-hACE2 mice with intense apical expression (E and F). ACE2 expression within the olfactory bulb (G-I) and the brain (J-L) was restricted to capillary endothelium with no neuronal expression. DAB, 200X total magnification. Bar = 100 μm.

We therefore aimed at analyzing expression and distribution of *hACE2* mRNA using RNAscope® ISH (Fig.12). Although no expression of *hACE2* mRNA was detected in the lungs of non-transgenic C57BL/6J mice (Fig. 12A), expression of *hACE2* mRNA was detectable, but of low expression in the lungs of K18-hACE2 mice, and mostly involved bronchiolar epithelial cells with sporadic expression in AT2 pneumocytes (Fig. 12B,C). These findings therefore suggest that *hACE2* expression might not be the sole host factor determinant of susceptibility to SARS-CoV-2. This is clearly exemplified by the following: 1) certain cell types that, while expressing *hACE2,* were non-permissive to SARS-CoV-2 infection throughout the experiment (i.e. bronchiolar epithelial cells); and 2) the near diffuse infection of AT1 cells by 4 dpi despite absent expression of *hACE2* in these cells. Altogether, these observations then support evidence for an ACE2-independent viral entry mechanism playing a major role in the pulmonary dissemination of K18-hACE2 mice.

**Fig. 12.**
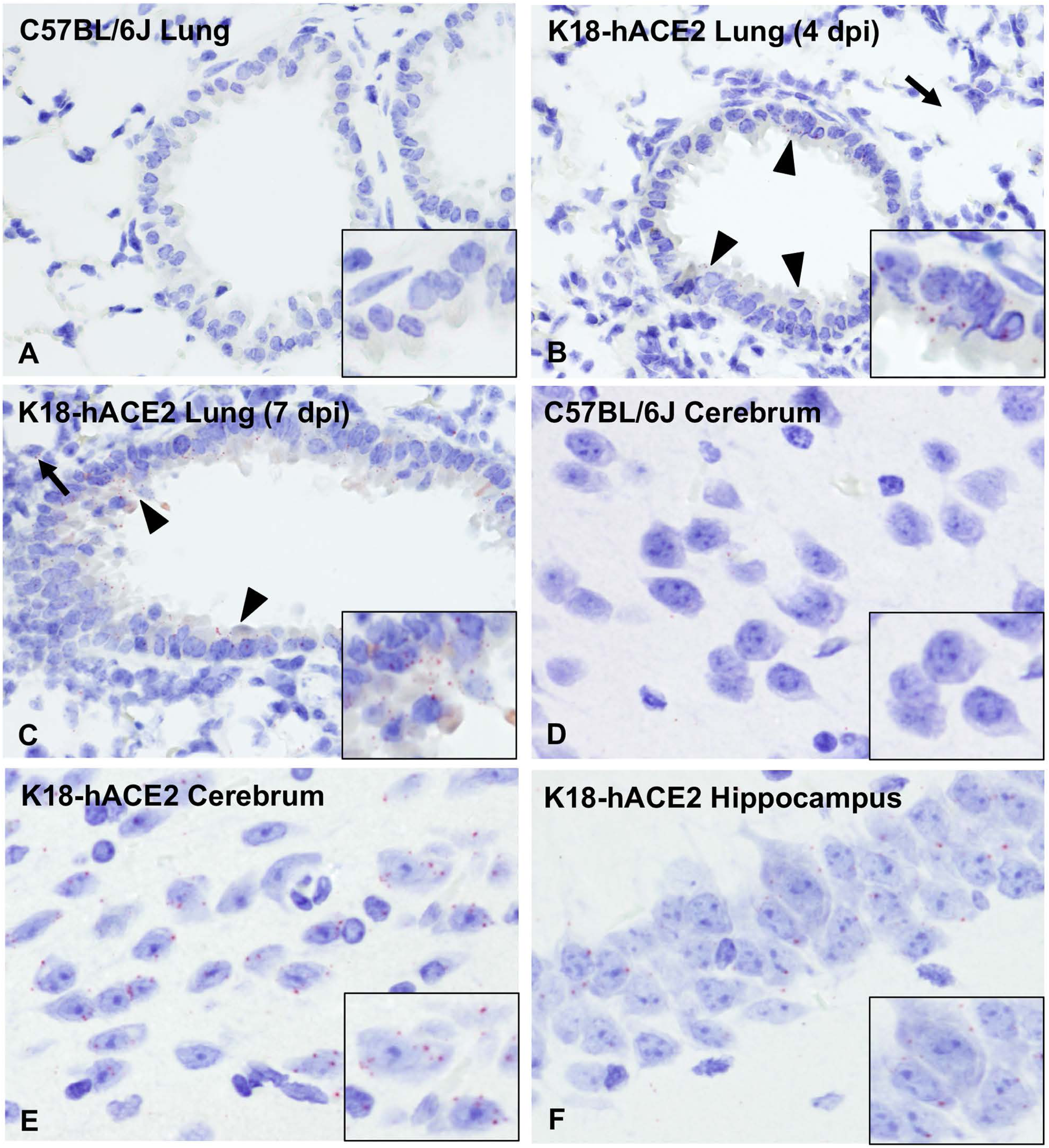
Expression and distribution of *hACE2* mRNA in the brain and lungs of C57BL/6J and K18-hACE2 transgenic mice via RNAscope® ISH. (A-C) *hACE2* lung expression. While no expression of *hACE2* was noted in the lungs of wild-type C57BL/6J mice (A), *hACE2* was expressed in the bronchiolar epithelium (arrowheads) and sporadic AT2 cells (arrows) in transgenic K18-hACE2 mice (B and C), which correlated with immunohistochemical findings. (D-F) *hACE2* brain expression. *hACE2* was not expressed in the Cerebrum of C57BL/6J mice (D) but in clusters of neurons within the cerebrum (E) and hippocampus (F). Fast Red, 400X total magnification. Bar = 50 μm.

In contrast to the lung, ACE2 protein was clearly overexpressed in the nasal cavity of K18-hACE2 mice compared to C57BL/6J mice. We assessed ACE2 protein expression on the rostral transitional epithelium, respiratory epithelium at the level of the intermediate turbinates, as well as in the ONE and olfactory bulb (Fig. 11D-F, G-I). Unlike C57BL/6J mice, in which ACE2 was undetectable within the nasal cavity, ACE2 protein was diffusely expressed within the apical membrane of transitional and respiratory epithelium, and segmentally within the apical surface of the ONE in both sham-inoculated and SARS-CoV-2-infected K18-hACE mice (Fig. 11D-F). For the olfactory bulb, olfactory neuroepithelium and respiratory epithelium of rostral turbinates, estimation of *hACE2* abundance and distribution could not be accurately assessed since the decalcification procedure is believed to have had a significant impact in the quality of cellular mRNA as demonstrated by the low detection of the housekeeping mRNA, *Ppib* (data not shown).

In the brain of both C57BL/6J and K18-hACE2 mice, ACE2 protein was observed in the vascular endothelium lining blood vessels (Fig. 11J-L), as well as ependymal and choroid plexus epithelium. In contrast, distribution of *hACE2* mRNA expression involved clusters of neurons within the cerebral cortex, hippocampus, midbrain, brainstem, and Purkinje cells from the cerebellum, with no expression noted in non-transgenic C57BL/6J mice (Fig. 12D-F). There was no expression of *hACE2* mRNA in vascular endothelial cells. Taken together, our data show a discrepancy between ACE2 protein and RNA expression and distribution within the CNS. This is partly attributable to the fact that the ACE2 antibody we utilized cross reacts with both hACE2 and mACE2 proteins, while the *ACE2* probe employed was human specific. The absence of *hACE2* hybridization with simultaneous ACE2 immunoreactivity in the capillary endothelium supports the notion that ACE2 expression in these cells is of murine origin. The absence of ACE2 immunoreactivity in neurons is suggestive of a potential restriction in the translation (or post-translation) of the ACE2 protein in these cells. This, in addition to the fact that Purkinje cells of the cerebellum do not appear permissive to SARS-CoV-2 infection despite the low expression of *hACE2* mRNA, suggests that ACE2 is likely not the sole host factor associated with neuroinvasion and that other ACE2-independent entry mechanisms contribute to neuroinvasion and spread by SARS-CoV-2 in this murine model. Alternatively, and/or in parallel, the overexpression of ACE2 protein within the nasal passages may be sufficient to enhance neuroinvasion by enhancing axonal transport via the ONE.

### Absence of infection and histologic lesions in extrapulmonary and extraneural tissues despite ACE2 expression

Other tissues examined included heart, kidney, stomach, duodenum, jejunum, ileum, cecum, and colon. All of these were histologically within normal limits (data not shown). No SARS-CoV-2 S protein was detected in any of these tissues at any time point (Table 1). ACE2 distribution was evaluated in sections of the heart, stomach, small intestine, and colon. While ACE2 expression was limited to the capillary vascular endothelium in the heart and glandular stomach, intense expression was noted in the non-glandular mucosa of the stomach (Fig.13 A-C) and apical surface of enterocytes lining the small intestinal mucosa (Fig.13 D-F). Colonic enterocytes rarely expressed ACE2 (Fig. 13G-I).

**Fig. 13.**
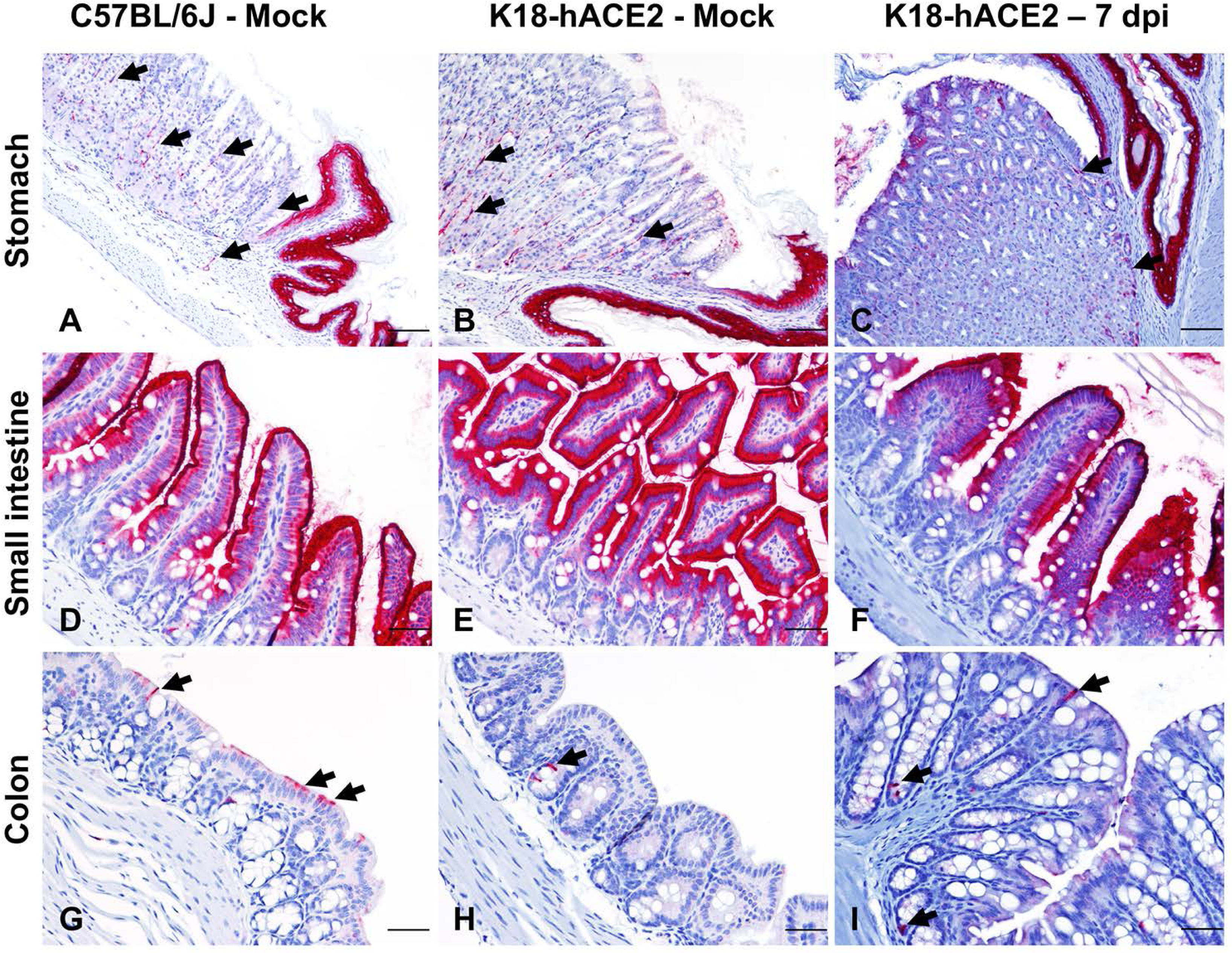
Expression of ACE2 in the gastrointestinal tract. Immunohistochemistry was performed using a cross-reactive anti-ACE2 antibody in the stomach (A-C), small intestine (D-F) and colon (G-I). In the stomach, ACE2 expression (red) was intense in the non-glandular mucosa and capillaries of the glandular mucosa (A-C, arrows). Enterocytes lining the small intestine of C57BL/6J and K18-hACE2 mice displayed prominent apical cytoplasmic ACE2 expression (D-F). In the colon, scattered enterocytes expressed ACE2 (G-I, arrows). Fast Red, 100X (A-C; bar = 200 μm) and 200X (D-I; bar = 100 μm) total magnification.

## DISCUSSION

The K18-hACE2 transgenic mouse model has become a widespread laboratory animal model suitable for studying SARS-CoV-2 pathogenesis as well as medical countermeasures against COVID-19 (20). The suitability of this model relies on the common host entry receptor shared between SARS-CoV and SARS-CoV-2 (20, 39), and transgenic mice expressing hACE2 under the K18 promoter develop lethal clinical disease associated with pulmonary pathology and neuroinvasion, with high viral titers (30, 36–38, 40, 47–49). In contrast, other murine models of SARS-CoV-2 (e.g. adenovirus-transduced hACE2 mice and hACE2 knock-in mice) develop only mild disease with limited and short-lived viral replication and pulmonary pathology, and low to no lethality (37, 50). While the K18-hACE2 murine model has been critical in shedding light on mechanisms of lung injury and dysfunction, it fails to faithfully recapitulate several key histologic features of severe and lethal cases of COVID-19 in humans, such as diffuse alveolar damage (DAD) with hyaline membrane formation and multi-organ failure associated with hypercoagulability and widespread microthrombi formation (43, 51).

To better understand the pathogenesis of SARS-CoV-2, well-characterized animal models are critically needed (22). Even though the K18-hACE2 murine model is currently under extensive use, several aspects associated with the temporospatial dynamics of SARS-CoV-2 infection remain poorly characterized, including the expression and cellular distribution of hACE2. In this work we further characterized pathological aspects related to viral pathogenesis in this unique murine model and hypothesized that the temporospatial distribution of SARS-CoV-2 and pathological outcomes following infection in the K18-hACE2 murine model is partially but not solely associated with hACE2 and that increased lethality in this model is related to neuroinvasion. The study presented herein provides additional novel information regarding the temporal and spatial aspects of SARS-CoV-2 infection in the K18-hACE2 mouse model with emphasis on pathological outcomes as well as a thorough and methodical characterization of ACE2 expression in this transgenic mouse model, which contributes to our understanding of this critical model used for preclinical evaluation of vaccines and antiviral therapeutics. Our findings not only demonstrate that lethality of this murine model is associated with neuroinvasion and subsequent neuronal cytopathic effect, but that SARS-CoV-2 tropism is not solely restricted to ACE2-expressing cells in K18-hACE2 mice. Thus, the neuropathogenic potential of SARS-CoV-2 is dependent on other currently unknown host factors.

Herein, we utilized a large cohort of K18-hACE2 mice enrolled in either a 14-day natural history or pre-determined serial euthanasia study to sequentially evaluate SARS-CoV-2 tropism and pathological alterations, spatial and temporal analysis of host factors including inflammatory response and ACE2/*hACE2* expression, and several clinical indices. Survival curve analysis demonstrated that lethality in infected mice only occurs at or after 6 dpi, and in most mice (∼94%), coincided with the initiation of neurologic signs and/or symptoms, neuronal cytopathic effect, and abundance of viral S protein, RNA, and infectious viral particles in the CNS. These observations clearly indicate neuroinvasion as a key determinant in the fatal outcome affiliated with this model. Our study also demonstrates that SARS-CoV-2 has a tropism for neurons within the spinal cord (predominantly within the cervicothoracic segments), which was only observed in mice with severe concurrent brain involvement. This observation could reflect descending progression originating from the brain, or alternatively axonal transport via motor or sympathetic sensory fibers. Concurrent brain and spinal cord disease rationalize the neurologic signs observed with this model, which included decreased mobility/responsiveness and decreased urine voiding, reflective of severe urinary bladder dilation and accumulation of concentrated urine. Given the spinal cord involvement, the latter is potentially attributed to altered spinal reflexes and/or decreased intervention of the detrusor muscle, which is required for normal micturition. An additional striking clinical feature in infected K18-hACE2 mice at 7 dpi was hypothermia, which is likely a consequence of hypothalamic (controls thermoregulation) and generalized neuronal dysfunction associated with SARS-CoV-2 neurotropism and serves as a clear clinical indicator of CNS involvement in this model. Our results unequivocally demonstrate that neuroinvasion is a major driver of fatality in this animal model compared to others such as Syrian hamsters, which display more severe pulmonary disease and infection of the ONE but lack evidence of neuroinvasion (52). Furthermore, these animals invariably recover within 14 days following intranasal infection with SARS-CoV-2 (24, 29, 52–54). Very few infected K18-hACE2 mice (2/30) from our survival curve study (14 dpi) survived and, while residual pulmonary inflammation was observed, these animals did not exhibit any evidence of neuroinvasion. Uniquely, both survivors developed pulmonary interstitial aggregates of B lymphocytes which were not observed at earlier time points, suggestive of the development of protective adaptive humoral response further supported by presence of neutralizing antibodies in these two animals, when compared to naïve animals. The absence of any overt neurological clinical signs, normal histologic appearance of the CNS and, absence of detectable SARS-CoV-2 protein or RNA in the two surviving mice supports the notion that animals can either fully recover from a milder form of neuroinvasion, or more likely, that in rare instances neuroinvasion fails to occur for unknown reasoning. Furthermore, we acknowledge that extensive neurobehavior testing, which is beyond the expertise of the authors, would be required to rule out any long-term neurological sequelae in the rare instance of survivors. Overall, these findings are of importance to researchers with a particular interest in studying SARS-CoV-2- associated neuropathogenesis, as premature euthanasia due to other clinical features (i.e., weight loss, ruffled fur, and/or respiratory distress) have the potential to precede CNS disease. Such terminal endpoints, if elected, may preclude evaluation of the effects of SARS-CoV-2 in the CNS. Instead, decreased responsiveness/mobility, tremors, and hypothermia should be interpreted to reflect better clinical findings supportive of neuroinvasive disease.

To date, the precise mechanism(s) enabling neuroinvasion in the K18-hACE2 model is poorly understood (11, 13, 15, 16, 52). Here, we determined that K18-hACE2 transgenic mice show a significant upregulation in the expression of ACE2 in the nasal cavity compared to wild-type C57BL/6J mice, in which ACE2 expression is undetectable by IHC. This difference between K18-hACE2 and C57BL/6J mice is clearly attributed to the expression of the *hACE2* transgene and is a key feature to the neuropathogenesis of this model. Interestingly, temporal analysis of SARS-CoV-2 S protein and RNA in the ONE of transgenic mice preceded and/or occurred simultaneously with infection of neurons within the glomerular and mitral layers of the olfactory bulb, supporting axonal transport through the cribriform plate as a primary portal of neuroinvasion. Expression of *hACE2* within neurons in the CNS is overall low and does not directly correlate with our immunohistochemical findings, where ACE2 protein was restricted to capillary endothelium, ependymal and choroid epithelium with sparing of neurons and their processes. These findings suggest the ACE2 expression in these anatomical compartments could be attributed to *mACE2* and/or indicative of a post-transcriptional event that could be limiting neuronal expression of *hACE2*. These along with the fact that *hACE2* mRNA is not abundantly and equally expressed among different neuronal populations and that Purkinje cells in the cerebellum express *hACE2* mRNA but are not permissive to SARS-CoV-2 infection, suggest that entry of SARS-CoV-2 into neurons is likely mediated by other host receptors independent of ACE2. Alternatively, overexpression at the interface of the ONE and neuronal synapses may be sufficient to rationalize the severe neuroinvasion observed in this model.

Infection of brain organoids has been shown to be inhibited using anti-ACE2 antibodies (19). However, brain organoids do not recapitulate the complexity of the entire CNS, and axonal transport of viral particles into the CNS can hardly be modeled *in vitro*. Altogether, this suggests that while ACE2 is assuredly an important mediator of CNS neuroinvasion, studying mechanisms of SARS-CoV-2 neuroinvasion likely require the use of more complex experimental systems. Neuropilin-1, a transmembrane glycoprotein serving as cell surface receptor for semaphorins and other ligands, as well as Tetraspanin 8 (TSPAN8), have recently been proposed as alternative host receptors for SARS-CoV-2 entry (55, 56). Even though we analyzed the expression of neuropilin-1 in this study (data not shown), we observed ubiquitous expression in the nasal passages, brain, kidneys, liver, and lungs, precluding any definitive conclusions in support or against these claims (55).

Anosmia and ageusia (loss of smell and taste, respectively) represent the earliest and most common but transient neurologic symptoms in people with COVID-19, being reported in ≥ 50% of cases (12, 13, 17). Hyposmia or anosmia has also been clearly characterized in K18-hACE2 mice, occurring between 2-3 dpi, which was characterized through a series of unique behavioral tests requiring a normal sense of smell (38). Other neurologic manifestations of COVID-19 have been attributed to acute cerebrovascular disease, with cohort studies reporting strokes in 2–6% of hospitalized patients (7, 13). Long-term neurologic sequelae associated with COVID-19 or its effect on neurodegenerative diseases remain unclear (7). Very little is known about the pathogenesis of these neurologic manifestations and whether they are directly or indirectly associated with SARS-CoV-2. ACE2 expression has been described in humans both in health and with chronic rhinosinusitis, with expression noted in sustentacular cells of the ONE, but not within immature and mature olfactory neurons (57). This observation led the authors to suggest that anosmia in COVID-19 is likely attributable to an indirect effect of SARS-CoV-2 infection. However, recent studies evaluating the brain and nasal autopsies from patients who died of COVID-19, detected SARS-CoV-2 protein and RNA in cells of neural origin within the ONE and cortical neurons occasionally associated with locally ischemic regions (18, 19). These studies provide evidence that the K18-hACE2 mice could have translational significance, even though ischemic lesions have not been reported including results from our study. Even though SARS-CoV-2 infects sustentacular cells within the neuroepithelium of Syrian hamsters (52), the K18-hACE2 and a transgenic mice expressing hACE2 under the HFH4 promoter are the only published models that consistently develop neuroinvasion with wild-type virus and, thus, will be particularly useful for studying SARS-CoV-2 neuropathogenesis, particularly the mechanisms of viral trafficking of into the CNS through the ONE (35).

Another important observation of the K18-hACE2 model is that SARS-CoV-2 tropism extensively involves infection of ACE2 and *hACE2* negative cells, including certain population of neurons and the vast majority of AT1 pneumocytes. Similarly, sole expression of *hACE2* in some cell types (i.e., CNS capillaries and bronchiolar epithelial cells) clearly does not render these cells susceptible to SARS-CoV-2 even following intranasal exposure and underscores the notion that other undetermined host factors are likely required to allow viral entry. Therefore, this model is relevant for investigating the role of alternative ACE2-independent entry mechanisms.

In conclusion, this study provides a comprehensive spatiotemporal analysis of SARS-CoV-2 infection in the K18-hACE2 transgenic murine model along with an analysis of the contribution of ACE2 in the permissiveness of the model. Our work provides extensive evidence that SARS-CoV-2 exhibits a marked neurotropism that is associated with lethality, and that this process likely occurs through mechanisms that are in part hACE2- independent. Although we documented significant reactive microgliosis and astrogliosis in terminal neuroinvasive disease, the exact role and molecular determinants of these observations, and their role in neuronal injury of the K18hACE2 model warrants further research. Lethal CNS invasion, combined with the absence of severe pulmonary hallmarks associated with lethal COVID-19, therefore calls for attentive caution when utilizing the K18-hACE2 mouse model to investigate certain aspects of SARS-CoV-2 pulmonary pathogenesis. Furthermore, due to the acute and fulminant neuroinvasion of this model, the protective ability of anti-viral therapies and T-cell based vaccines against lethal challenge in this model might indeed be underestimated, which is reflected in several studies that have utilized terminal timepoints proceeding neuroinvasion as their efficacy endpoints (58–60). Regardless, the K18-hACE2 mouse model represents a promising model for understanding the mechanisms governing SARS-CoV-2 neuroinvasion, ACE2-independent virus entry, and evaluating potent and fast-acting prophylactic countermeasures. Lastly, this model may serve useful in evaluating efficacy of therapeutics to block development of reactive/injurious microglial and/or astrocyte phenotypes if determined to play a key role in the neuronal injury observed in this model.

## MATERIALS AND METHODS

### Biosafety

All aspects of this study were approved by the Institutional Biosafety Committee and the office of Environmental Health and Safety at Boston University prior to study initiation. Work with SARS-CoV-2 was performed in a biosafety level-3 laboratory by personnel equipped with powered air-purifying respirators.

### Cells and viruses

African green monkey kidney Vero E6 cells (ATCC^®^ CRL-1586™, American Type Culture Collection, Manassas, VA) were maintained in Dulbecco’s minimum essential medium (DMEM; Gibco, Carlsbad, CA [#11995-065]) containing 10% fetal bovine serum (FBS, ThermoFisher Scientific, Waltham, MA), 1X non-essential amino acids (ThermoFisher Scientific), penicillin and streptomycin (100 U/ml and 100 μg/ml), and 0.25 μg/ml of amphotericin B (Gibco^®^, Carlsbad, CA), and incubated at 37 °C and 5% CO_2_ in a humidified incubator.

### SARS-CoV-2 isolate stock preparation and titration

All replication-competent SARS-CoV-2 experiments were performed in a biosafety level 3 laboratory (BSL-3) at the Boston University’ National Emerging Infectious Diseases Laboratories. 2019-nCoV/USA-WA1/2020 isolate (NCBI accession number: MN985325) of SARS-CoV-2 was obtained from the Centers for Disease Control and Prevention (Atlanta, GA) and BEI Resources (Manassas, VA). To generate the passage 1 (P1) virus stock, Vero E6 cells, pre-seeded the day before at a density of 10 million cells, were infected in T175 flasks with the master stock, diluted in 10 ml final volume of Opti-MEM (ThermoFisher Scientific). Following virus adsorption to the cells at 37 °C for 1 h, 15 ml DMEM containing 10% FBS and 1X penicillin/streptomycin was added to the flask. The next day, media was removed, the cell monolayer was rinsed with 1X phosphate buffered saline (PBS) pH 7.5 (ThermoFisher Scientific) and 25 ml of fresh DMEM containing 2% FBS was added. Two days later, when the cytopathic effect of the virus was clearly visible, culture medium was collected, filtered through a 0.2 µm filter, and stored at -80 °C. Our P2 working stock of the virus was prepared by infecting Vero E6 cells with the P1 stock, at a multiplicity of infection (MOI) of 0.1. Cell culture media was harvested at 2 and 3 dpi, and after the last harvest, ultracentrifuged (Beckman Coulter Optima L-100k; SW32 Ti rotor) for 2 h at 25,000 rpm (80,000 *X* g) over a 20% sucrose cushion (Sigma-Aldrich, St. Louis, MO). Following centrifugation, the media and sucrose were discarded, and pellets were left to dry for 5 minutes at room temperature. Pellets were then resuspended overnight at 4 °C in 500 µl of 1X PBS. The next day, concentrated virions were aliquoted at stored at -80 °C.

The titer of our viral stock was determined by plaque assay. Vero E6 cells were seeded into a 12-well plate at a density of 2.5 x 10^5^ cells per well and infected the next day with serial 10-fold dilutions of the virus stock for 1 h at 37 °C. Following virus adsorption, 1 ml of overlay media, consisting of 2X DMEM supplemented with 4% FBS and mixed at a 1:1 ratio with 1.2% Avicel (DuPont; RC-581), was added in each well. Three days later, the overlay medium was removed, the cell monolayer was washed with 1X PBS and fixed for 30 minutes at room temperature with 4% paraformaldehyde (Sigma-Aldrich). Fixed cells were then washed with 1X PBS and stained for 1h at room temperature with 0.1% crystal violet (Sigma-Aldrich) prepared in 10% ethanol/water. After rinsing with tap water, the number of plaques were counted, and the virus titer was calculated. The titer of our P2 virus stock was 4 x 10^8^ plaque forming units (PFU)/ml.

### Recombinant SARS-CoV-2 NanoLuciferase stock

Recombinant SARS-CoV-2 virus expressing a NanoLuciferase reporter (rSARS-CoV-2 NL) (61) was generously provided by the Laboratory of Pei-Yong Shi. A day prior to propagation 10 million Vero E6 cells were seeded in a T-175 flask. To grow virus, 10 µl of rSARS-CoV-2 NL virus stock was diluted in 10 ml of OptiMEM media (ThermoFisher Scientific, #51985091) and then added to cells. Virus was incubated on cells for 1 hour at 37°C then 15 mL of DMEM containing 10% FBS and 1% penicillin/streptomycin was added. The morning after infection, media was removed, cells were washed once with 1X PBS and 25 ml of fresh DMEM containing 2% FBS and 1% penicillin/streptomycin was added to the flask. Virus was incubated for an additional 48 hours, supernatant was collected, filtered through a 0.22 µM filter, and stored at -80°C. To concentrate virus, the stock was thawed and concentrated by ultracentrifugation (Beckman Coulter Optima L-100k; SW32 Ti rotor) at 25,000 x g for 2 hours at 4 °C on a 20% sucrose cushion (Sigma-Aldrich, St. Louis, MO). Media and sucrose were decanted, pellets were allowed to dry for 5 minutes at room temperature, then viral pellets were suspended in 100 µl of 1X PBS and left at 4 °C overnight. The next day, concentrated virus was aliquoted and stored at -80 °C.

### Mice

Mice were maintained in a facility accredited by the Association for the Assessment and Accreditation of Laboratory Animal Care (AAALAC). All protocols were approved by the Boston University Institutional Animal Care and Use Committee (PROTO202000020). Heterozygous K18-hACE2 C57BL/6J mice of both sexes (strain: 2B6.Cg-Tg(K18-ACE2)2Prlmn/J) were obtained from the Jackson Laboratory (Jax, Bar Harbor, ME). Animals were group-housed by sex in Tecniplast green line individually ventilated cages (Tecniplast, Buguggiate, Italy). Mice were maintained on a 12:12 light cycle at 30-70% humidity and provided ad-libitum water and standard chow diets (LabDiet, St. Louis, MO).

### Intranasal inoculation with SARS-CoV-2

At 4 months of age, K18-hACE2 mice of both sexes were intranasally inoculated with 1 x 10^6^ PFU of SARS-CoV-2 in 50 µl of sterile 1X PBS (n=61 [n=34 male and n=27 female], or sham inoculated with 50 µl of sterile 1X PBS (n=3; female). Inoculations were performed under 1-3% isoflurane anesthesia. Thirty-eight of these animals were enrolled in a 14-day survival curve study. For histologic analysis, twenty-six animals were examined (n=15 male and n=11 female), which included three female Sham/PBS inoculated controls and predetermined euthanasia timepoints at 2 and 4 dpi prior to animals reaching euthanasia criteria.

### Clinical monitoring

Animals included in the 14-day survival curve study were intraperitoneally implanted with an RFID temperature-monitoring microchip (Unified Information Devices, Lake Villa, IL, USA) 48-72 hours prior to inoculation. An IACUC-approved clinical scoring system was utilized to monitor disease progression and establish humane endpoints (Table 2). Categories evaluated included body weight, general appearance, responsiveness, respiration, and neurological signs for a maximum score of 5. Animals were considered moribund and humanely euthanized in the event of the following: a score of 4 or greater for 2 consecutive observation periods, weight loss greater than or equal to 20%, severe respiratory distress, or lack of responsiveness. Clinical signs and body temperature were recorded once per day for the duration of the study. For design of the survival curve, animals euthanized on a given day were counted dead the day after. Animals found dead in cage were counted dead on the same day.

**Table 2.** Clinical scoring system used for clinical monitoring of SARS-CoV-2-infected K18-hACE2 mice.

| Category | Score = Criteria |
| --- | --- |
| Body weight | 1 = 10-19% loss |
| Respiration | 1 = rapid, shallow, increased effort |
| Appearance | 1 = ruffled fur, hunched posture |
| Responsiveness | 1 = low to moderate unresponsiveness |
| Neurologic signs | 1 = tremors |

**Table 3.** Antibodies and antigen retrieval conditions for the assays performed in this study.

|  | Sequence | Antigen Target | Species Origin | Clone | Manufacturer | Catalog | Primary Antibody Dilution | Antigen Retrieval (Ventana) | Chromogen or Fluorophore |
| --- | --- | --- | --- | --- | --- | --- | --- | --- | --- |
| Assay 1 | NA | SARS-CoV-2 Spike (S) | Mouse | E7U6O | Cell Signaling Technology | Pre-commercialization | 1:1000 | CC1 (Tris) | DAB |
| Assay 2 | NA | Angiotensin converting enzyme 2 (ACE2) | Rabbit | EPR34435 | Abcam | ab108252 | 1:200 | CC1 (Tris) | Discovery Red and DAB |
| Assay 3 | 1 | SARS-CoV-2 S | Mouse | E7U6O | Cell Signaling Technology | Pre-commercialization | 1:1000 | CC1 (Tris) | Opal 480 |
|  | 2 | Iba-1 | Rabbit | Polyclonal | WAKO | 019-19741 | 1:2000 | CC2 (Citrate) | Opal 570 |
|  | 3 | GFAP | Rabbit | Polyclonal | DAKO | Z0334 | 1:500 | CC1 (Tris) | Opal 690 |
| Assay 4 | 1 | CD8 | Rabbit | D4W2Z | Cell Signaling Technology | 98941 | 1:200 | CC1 (Tris) | Opal 620 |
|  | 2 | SARS-CoV-2 S | Mouse | E7U6O | Cell Signaling Technology | Pre-commercialization | 1:1000 | CC1 (Tris) | Opal 570 |
|  | 3 | CD19 | Rabbit | D4V4B | Cell Signaling Technology | 90176 | 1:600 | CC2 (Citrate) | Opal 520 |
|  | 4 | Iba-1 | Rabbit | Polyclonal | WAKO | 019-19741 | 1:2000 | CC2 (Citrate) | Opal 690 |
| Assay 5 | 1 | RAGE | Rat | EPR21171 | R&D | MAB1179q-100 | 1:50 | CC1 (Tris) | Opal 480 |
|  | 2 | SARS-CoV N | Rabbit | Polyclonal | Novus biologicals | NB100-56576 | 1:200 | CC1 (Tris) | Opal 570 |
|  | 3 | Prosurfactant C Protein | Rabbit | Polyclonal | Seven Hills Bioreagents | WRAB-9337 | 1:800 | CC2 (Citrate) | Opal 690 |
|  | 4 | CD31 | Rabbit | D8V9E | Cell Signaling Technology | 77699S | 1:100 | CC2 (Citrate) | Opal 520 |

### In vivo 3D-imaging and analysis

K18-hACE2 mice were infected with 1×10^6 PFU of SARS-CoV-2 NL in 50 μl of 1X PBS via intranasal inoculation. To image, mice were administrated two, 75 μl subcutaneous injections of 1X PBS containing 0.65 uM Fluorofurimazine (FFz) substrate (Promega) for a total of 1.3 uM FFz per mouse. Mice were then imaged using a 3D-imaging mirror gantry isolation chamber (InVivo Analytics) and an IVIS spectrum imager (PerkinElmer). To perform imaging, mice were anesthetized with 2.5% isoflurane, placed into a body conforming animal mold (BCAM) (InVivo Analytics), and then imaged within 5 minutes of FFz injection. Images were acquired using a sequence imaging as followed; 60 seconds (s) open filter, 240 s 600 nm, 60 s open, 240 s 620 nm, 60 s open, 240 s 640 nm, 60 s open, 240 s 660 nm, 60 s open, 680 nm, 60 s open. Data analysis was performed using the cloud-based InVivoPlot software (InVivo Analytics).

### Tissue processing and viral RNA isolation

Tissues were collected from mice and stored in 600 µl of RNA*later* (Sigma-Aldrich; # R0901500ML) and stored at -80 °C. For processing, 20 – 30 mg of tissue were placed into a 2 ml tube with 600 µl of RLT buffer with 1% β-mercaptoethanol and a 5 mm stainless steel bead (Qiagen, Valencia, CA; #69989). Tissues were then dissociated using a Qiagen TissueLyser II (Qiagen) with the following cycle parameters: 20 cycles/s for 2 min, 1 min wait, 20 cycles/s for 2 min. Samples were centrifuged at 17,000 *X* g (13,000 rpm) for 10 minutes and supernatant was transferred to a new 1.5 ml tube. Viral RNA isolation was performed using a Qiagen RNeasy Plus Mini Kit (Qiagen; #74134), according to the manufacturer’s instructions, with an additional on-column DNase treatment (Qiagen; #79256). RNA was finally eluted in 30 μl of RNase/DNase-free water and stored at -80 °C until used.

### Quantification of infectious particles by plaque assay

Quantification of SARS-CoV-2 infectious particles were quantified by plaque assay. After euthanizing mice, tissues were collected in 600 µL of RNA*later* (ThermoFisher Scientific, AM7021) and stored at - 80 C until analysis. The day prior to experiments, 24-well plates containing 8×104 Very E6 cells per well were plated. Between 20-40 mg of tissue was weighed out and placed into a 2 ml tube containing 500 μl of OptiMEM (ThermoFisher) and a 5mm Steal Bead (Qiagen #69997). Tissues were then homogenized using a Qiagen TissueLyser II (Qiagen; Germantown, MD) by two dissociations cycles (two-minutes at 1,800 oscillations/minute) with a one-minute rest in between. Samples were then subject to centrifugation with a benchtop centrifuge at 13,000 rpm for 10 minutes and supernatant was transferred to a new 1.5 ml tube. From this, 1:10 – 1:10^6^ dilutions were made in OptiMEM and 200 µl of each dilution were plated onto 24-well plates. Media was incubated at 37 °C for 1 hour with gentle rocking of the plate every 10 minutes. After viral adsorption, 800 μl of a 1:1 mixture of 2X DMEM containing 4% FBS 1% penicillin/streptomycin and 2.4% Avicel (Dupont) was overlaid into each well. Cells were then incubated for 72 hours at 37°C with 5% CO_2_. After incubation, Avicel was removed, cells were washed with 1X PBS, and cells were fixed in 10% formalin for 1 hour. After fixation, formalin was removed, cells were stained with 0.1% crystal violet in 10% ethanol/water for 30 minutes and washed with tap water. Plates were then dried, the number of plaques were counted, and infectious particles (PFU/mg of tissue) were calculated.

### RNA isolation from serum

Total viral RNA was isolated from serum using a Zymo Research Corporation Quick-RNA^TM^ Viral Kit (Zymo Research, Tustin, CA; #R1040) according to the manufacturer’s instructions. RNA was eluted in 15 μl of RNase/DNase-free water and stored at -80 °C until used.

### SARS-CoV-2 E-specific reverse transcription quantitative polymerase chain reaction (RT-qPCR)

Viral RNA was quantitated using single-step RT-quantitative real-time PCR (Quanta qScript One-Step RT-qPCR Kit, QuantaBio, Beverly, MA; VWR; #76047-082) with primers and TaqMan® probes targeting the SARS-CoV-2 E gene as previously described (62). Briefly, a 20 μl reaction mixture containing 10 μl of Quanta qScript™ XLT One-Step RT-qPCR ToughMix, 0.5 μM Primer E_Sarbeco_F1 (ACAGGTACGTTAATAGTTAATAGCGT), 0.5 μM Primer E_Sarbeco_R2 (ATATTGCAGCAGTACGCACACA), 0.25 μM Probe E_Sarbeco_P1 (FAM-ACACTAGCCATCCTTACTGCGCTTCG-BHQ1), and 2 μl of template RNA was prepared. RT-qPCR was performed using an Applied Biosystems QuantStudio 3 (ThermoFisher Scientific) and the following cycling conditions: reverse transcription for 10 minutes at 55 °C, an activation step at 94 °C for 3 min followed by 45 cycles of denaturation at 94 °C for 15 seconds and combined annealing/extension at 58 °C for 30 seconds. Ct values were determined using QuantStudio^TM^ Design and Analysis software V1.5.1 (ThermoFisher Scientific). For absolute quantitation of viral RNA, a 389 bp fragment from the SARS-CoV-2 E gene was cloned onto pIDTBlue plasmid under an SP6 promoter using NEB PCR cloning kit (New England Biosciences, Ipswich, MA). The cloned fragment was then *in vitro* transcribed (mMessage mMachine SP6 transcription kit; ThermoFisher) to generate an RT-qPCR standard.

### Serum infectivity assay

One day prior to the experiment, 5×10^4^ Vero E6 cells were plated into a 24-well plate. Cells were then dosed with 200 μl of OptiMEM containing 20 μl of serum or SARS-CoV-2 WA-isolate (MOI=0.001 [positive control]), incubated for 1 hour at 37°C, media was removed and fresh DMEM containing 2% FBS and 1% penicillin/streptomycin was added. Cells were incubated at 37°C with 5% CO2 for 48 hours, 100 uL of supernatant was collected and RNA was extracted using a Quick-RNA Viral Kit as per manufacturer’ instructions (Zymo Research) for analysis by RT-qPCR.

### Serum neutralization assay

One day prior to the experiment, 1×10^4^ VeroE6 cells were plated into a 96-well plate. Serum was decomplemented at 56°C for 30 minutes. Serum was diluted 1:10 in OptiMEM and then serial diluted 2-fold for a total of ten-dilutions. Serum dilutions were mixed with SARS-CoV-2 NL virus (MOI=1), incubated for 1 hour at room temperature and then plated onto cells. After a 1-hour incubation at 37°C inoculum was removed and 200 μl of fresh DMEM containing 2% FBS and 1% penicillin/streptomycin was added. After a 24h incubation at 37°C with 5% CO2 media was removed and cells were fixed with 10% formalin for 1 hour. A SARS-CoV-2 spike neutralizing antibody (Sino Biological Inc.; 2ug/uL) was used as a positive control for neutralization. After fixation formalin was removed, cells were washed with 1X PBS and 20 uM furimazine (MedChem Express) luciferin substrate was added onto cells. Cells were then imaged using an IVIS spectrum imager (PerkinElmer) and analyzed using LivingImage software (PerkinElmer).

### Histology

Animals were anesthetized with 1-3% isoflurane and euthanized with an intraperitoneal overdose of ketamine and xylazine before harvest and fixation of tissues. Lungs were insufflated with ∼1.5mL of 1% low melting point agarose (Sigma-Aldrich) diluted in 1X PBS using a 24-gauge catheter placed into the trachea. The skull cap was removed and the animal decapitated and immersed in 10% neutral buffered formalin.

Additional tissues harvested included the heart, kidneys, and representative sections of the gastrointestinal tract, which included the duodenum, jejunum, ileum, cecum, and colon. Tissues were inactivated in 10% neutral buffered formalin at a 20:1 fixative to tissue ratio for a minimum of 72 hours before removal from BSL-3 in accordance with an approved institutional standard operating procedure. Following fixation, the whole head was decalcified in Immunocal™ Decalcifier (StatLab, McKinney, TX) for 7 days before performing a mid-sagittal section dividing the two hemispheres into even sections. Tissues were subsequently processed and embedded in paraffin following standard histological procedures. Five-micron sections were obtained and stained with hematoxylin and eosin or Luxol Fast Blue (myelin stain).

### Immunohistochemistry and RNAscope^®^ *in situ* hybridization

Immunohistochemistry (IHC) was performed using a Ventana BenchMark Discovery Ultra autostainer (Roche Diagnostics, Indianapolis, IN). Specific IHC assay details including antibodies, protein retrieval, sequence of multiplex assays, and incubation periods are found in Table 2 SARS-CoV-2 S was semiquantitatively scored as follows: 0, no viral protein observed; 1, up to 5% positive cells per 400X field examined; 2, 5-25% positive cells per 400X field examined; and 3, up to 50% positive cells per 400X field examined.

For SARS-CoV-2 RNAscope^®^ ISH, an anti-sense probe targeting the spike (S; nucleotide sequence: 21,563-25,384) of SARS-CoV-2, USA-WA1/2020 isolate (GenBank accession number MN985325.1) was used as previously described (23, 42). The RNAscope^®^ ISH assay was performed using the RNAscope 2.5 LSx Reagent Kit (Advanced Cell Diagnostics, Newark, CA) on the automated BOND RXm platform (Leica Biosystems, Buffalo Grove, IL) as described previously (23). Briefly, four-micron sections of formalin-fixed paraffin-embedded (FFPE) tissue was subjected to automated baking and deparaffinization followed by heat-induced epitope retrieval (HIER) using a ready-to-use EDTA-based solution (pH 9.0; Leica Biosystems) at 100 °C for 15 min. Subsequently, tissue sections were treated with a ready-to-use protease (RNAscope® 2.5 LSx Protease) for 15 min at 40 °C followed by a ready-to-use hydrogen peroxide solution for 10 min at room temperature. Slides were then incubated with the ready-to-use probe mixture for 2 h at 40 °C, and the signal amplified using a specific set of amplifiers (AMP1 through AMP6 as recommended by the manufacturer). The signal was detected using a Fast-Red solution for 10 minutes at room temperature. Slides were counterstained with a ready-to-use hematoxylin for 5 min, followed by five washes with 1X BOND Wash Solution (Leica Biosystems) for bluing. Slides were finally rinsed in deionized water, dried in a 60°C oven for 30 min, and mounted with Ecomount^®^ (Biocare, Concord, CA, USA). A SARS-CoV-2-infected Vero E6 cell pellet was used as a positive assay control. For all assays, an uninfected mouse was used as a negative control.

For *hACE2* mRNA RNAscope^®^ ISH, an anti-sense probe targeting *hACE2* (GenBank accession number NM_021804.3; Cat. No. 848038) with no cross-reactivity to murine *Ace2* was used in a similar manner as described above with the exception that AMP5 and AMP6 were incubated for 45 min and 30 min, respectively. Murine *peptidylprolyl isomerase B (Ppib)* mRNA was used as a housekeeping gene to determine RNA quality and a Vero E6 cell pellet was used as a positive assay control.

### Multispectral microscopy

Fluorescently labeled slides were imaged using a Mantra 2.0^TM^ or Vectra Polaris^TM^ Qunatitative Pathology Imaging System (Akoya Biosciences, Marlborough, MA). To maximize signal-to-noise ratios, images were spectrally unmixed using a synthetic library specific for the Opal fluorophores used for each assay and for 4′,6-diamidino-2-phenylindole (DAPI). An unstained lung or brain section were used to create a tissue specific autofluorescence signature that was subsequently removed from whole-slide images using InForm software version 2.4.8 (Akoya Biosciences).

### Quantitative Image analysis of multiplex immunohistochemistry

Digitized whole slide scans were analyzed using the image analysis software HALO (Indica Labs, Inc., Corrales, NM). Slides were manually annotated to include only the brain and/or lung parenchyma depending on the panel being evaluated. Visualization threshold values were adjusted in viewer settings to reduce background signal and fine-tune visibility of markers within each sample. For the CNS panel, area quantification (AQ) was performed to determine percentages of SARS-CoV-2 Spike, Iba1 (microglia) and GFAP (astrocyte) immunoreactivity. For the lung panel, we employed the HALO Highplex (HP) module which allows for simultaneous analysis of multiple fluorescent markers within a cellular compartment. Individual cells were identified using DAPI to segment individual nuclei. Minimum cytoplasm and membrane thresholds were set for each dye to detect positive staining within a cell. Parameters were set using the real-time tuning mechanism that was tailored for each individual sample based on signal intensity. Phenotypes were determined by selecting inclusion and exclusion parameters relating to stains of interest. We used the following phenotypes: CD8+ (cytotoxic T-cells), CD20+ (B-cells), and Iba1+ (macrophages). The algorithm produces a quantitative output for each cell phenotype as well as total cells per total area analyzed for an output of cells/µm^2^. The AQ module was also used the lung panel for quantification of SARS-CoV-2-Spike immunoreactivity.

### Quantitative image analysis of brightfield microscopy

Digitized whole slide scans of hematoxylin & eosin (H&E) stained k18 mouse lungs were analyzed using the Halo Tissue Classifier module. TC is a train-by-example machine learning algorithm used to identify dissimilar areas of tissue based on contextual features. For these lung samples, a classifier was created to distinguish areas of pneumonic lung from normal stroma. The classifier was run on whole lung images to determine the percentage of pneumonia. Quantitative outputs are given as total classified area (mm^2^), normal lung area (mm^2^), and pneumonia area (mm^2^). We divided pneumonic area by total classified area to generate a percentage of pneumonia for statistical analysis.

### Transmission electron microscopy

Tissue samples were fixed for 72 hours in a mixture of 2.5% Glutaraldehyde and 2% formaldehyde in 0.1 M sodium cacodylate buffer (pH 7.4). Samples were then washed in 0.1M cacodylate buffer and postfixed with 1% Osmiumtetroxide (OsO4)/1.5% Potassiumferrocyanide (KFeCN6) for 1 hour at room temperature. After washes in water and 50mM Maleate buffer pH 5.15 (MB), the samples were incubated in 1% uranyl acetate in MB for 1hr, washed in MB and water, and dehydrated in grades of alcohol (10min each; 50%, 70%, 90%, 2×10min 100%). The tissue samples were then put in propyleneoxide for 1 hr and infiltrated ON in a 1:1 mixture of propyleneoxide and TAAB Epon. The following day the samples were embedded in fresh TAAB Epon and polymerized at 60°C for 48 hrs. Semi-thin (0.5um) and ultrathin sections (50-80nm) were cut on a Reichert Ultracut-S microtome (Leica). Semi-thin sections were picked up on glass slides and stained with Toluidine blue for examination at the light microscope level to find affected areas in the tissue. Ultrathin sections from those areas were picked up onto formvar/carbon coated copper grids, stained with 0.2% lead citrate and examined in a JEOL 1200EX transmission electron microscope (JOEL, Akishima, Tokyo, Japan). Images were recorded with an AMT 2k CCD camera.

### Statistical analysis

Descriptive statistics and graphics as well as Kaplan-Meier (survival) curves and statistical tests were performed using GraphPad Prism v9.1.2 statistical analysis software (GraphPad, San Diego, CA). Clinical parameters and quantitative pathology results were analyzed using a one-way ANOVA with Dunnett post-hoc analysis with means of groups compared to the Sham-inoculated negative controls. Viral load data were evaluated using either a one-way (serum qPCR) or two-way ANOVA (tissue qPCR and PFU data) with Tukey post hoc analysis. Significance levels were set at p-value<0.05 in all cases. Statistical significance on figures and supplemental figures is labelled as follow: \**p*≤0.05, \*\**p*≤0.01, \*\*\**p*≤0.001, \*\*\*\**p*≤0.0001.

## ACKNOWLEDGEMENTS

We thank the Evans Center for Interdisciplinary Biomedical Research at Boston University School of Medicine for their support of the Affinity Research Collaborative on ‘Respiratory Viruses: A Focus on COVID-19’. This work utilized a Ventana Discovery Ultra autostainer that was purchased with funding from a National Institutes of Health SIG grant (S10-OD026983). This work was also supported by a Boston University Start-up fund, and a Peter Paul Career Development Professorship (to F.D.), as well as by grants from the National Institutes of Health (R21 ES032882, K22 AI144050 to F.D). This study was also partially supported by start-up funds provided by the School of Veterinary Medicine, Louisiana State University to Dr. Udeni Balasuriya (PG002165) and by pilot funding to Dr. Markus Bosmann from the National Institutes of Health grant 1UL1TR001430. Drs. Crossland and Carossino would like to thank our pathology mentors Drs. Fabio Del Piero and Ingeborg M. Langohr for helping instill our passion for pathology and for introducing us to each other. We are hopeful these efforts will represent the early days of a fruitful and long-lasting collaboration. We acknowledge the histology and immunohistochemistry sections at the Louisiana Animal Disease Diagnostic Laboratory for their technical assistance. The following reagent was deposited by the Centers for Disease Control and Prevention and obtained through BEI Resources, NIAID, NIH: SARS-Related Coronavirus 2, Isolate USA-WA1/2020, NR-52281. K.P.F reports that he is an employee of PerkinElmer, Inc., a manufacturer of diagnostic and analytical equipment. N.P. and A.K. declare the following competing interest as shareholders of InVivo Analytics with issued patents. T.A.K. and J.R.W. are both employees of Promega Corporation.

## AUTHOR CONTRIBUTIONS

F. Douam, NA. Crossland, M. Carossino, U. Balasuriya and M. Bosmann designed the study; F. Douam, NA. Crossland, M. Saeed, M. Carossino, P. Montanaro, A. O’Connell, D. Kenney, H. Gertje, K Grosz, Maria Ericsson, BR Huber and S Kurnick performed the experiments; F. Douam, N. Crossland, M. Carossino, P. Montanaro, A. O’Connell, and D. Kenney performed data analysis; T. Kirkland, J Walker, K Francis, and A Klose provided resources and visualization for *in vitro* imaging; and M. Carossino, NA. Crossland, and F. Douam wrote the original draft.

